# Regulation of lipid metabolism is a primordial function of STING

**DOI:** 10.1101/2025.11.17.688879

**Authors:** Soumyabrata Guha, Joanna Re, Morgane Chemarin, Stéphane Grégoire, Yasmine Messaoud-Nacer, Hanane Chamma, Adeline Augereau, Jennifer Barrat, Niyazi Acar, Isabelle K Vila, Pierre Boudinot, Nadine Laguette

## Abstract

The stimulator of interferon genes (STING) is a pivotal regulator of type I interferon (IFN) responses. Although the IFN system is confined to vertebrates, STING is present across metazoans and in some unicellular eukaryotes, suggesting involvement in distinct functions prior to vertebrate divergence. We here explored the conservation of STING-mediated regulation of polyunsaturated fatty acid (PUFA) metabolism. We found that tested STING homologs from vertebrates, invertebrates, and unicellular eukaryotes interacted with the fatty acid desaturase 2 (FADS2) rate-limiting enzyme in PUFA metabolism and subsequent functional outputs. The ability to regulate lipid metabolism did not correlate with antiviral activity, suggesting that the cooptation of this metabolic pathway by the IFN-based immune system is independent of STING-associated immune responses. Thus, STING-mediated metabolic regulation is an evolutionarily conserved feature and a primordial STING function.

## INTRODUCTION

The endoplasmic reticulum (ER)-resident stimulator of interferon genes (STING) adaptor protein is a central regulator of innate immune responses, initially identified as responsible for the induction of type I interferons (IFN) in response to cytosolic double-stranded deoxyribonucleic acid (dsDNA) in mammals(Barber, 2011; Ishikawa and Barber, 2008). This function of STING is initiated following the detection of exogenous and endogenous cytosolic dsDNA by the cyclic guanosine monophosphate (GMP)-adenosine monophosphate (AMP) (cGAMP) synthase (cGAS) pattern recognition receptor(Bregnard et al., 2016; Sun et al., 2013). Upon activation, cGAS catalyzes the production of the 2’3’-cGAMP cyclic dinucleotide (CDN), which subsequently binds to STING(Wu et al., 2013), inducing its oligomerization(Shang et al., 2019), translocation from the ER to the Golgi apparatus(Ishikawa et al., 2009), and assembly of the STING signalosome(Wu et al., 2013; Zhang et al., 2019). The latter canonically comprises the TANK binding kinase 1 (TBK1) that catalyzes the activating phosphorylation of STING(Zhang et al., 2019), as well as of transcription factors, such as interferon regulatory factors (IRF)(Chen and Jiang, 2013; McWhirter et al., 2004; Wu et al., 2013), culminating in the production of inflammatory cytokines and chemokines alongside type I IFNs(Ishikawa and Barber, 2008). Other well-documented immune-related functions of STING comprise the activation of the nuclear factor kappa B (NF-κB) signaling cascade(Abe and Barber, 2014), as well as triggering non-canonical autophagy(Gui et al., 2019; Prabakaran et al., 2018). Thus, STING activation is essential to shaping innate immune responses to dsDNA.

Analysis of the cGAS and STING encoding genes and of their immune-related functions in the choanoflagellate *Monosiga brevicollis* revealed that the eukaryotic cGAS-STING signaling pathway arose in unicellular eukaryotes over 600 million years ago, before the emergence of ancestral metazoans(Woznica et al., 2021). Indeed, STING homologs have been discovered in most metazoan phyla (with some exceptions such as nematodes), unicellular choanoflagellates, and even bacteria(Morehouse et al., 2020). However, neither its structure nor its canonical function, are strictly conserved across the tree of life. Indeed, to trigger pro-inflammatory signaling, the prototypical STING protein relies on three main domains(Wu et al., 2014): four N-terminal transmembrane (TM) domains ensure the anchoring of the protein to membranes, followed by a central CDN binding domain (CBD), and a C-terminal tail (CTT) that is essential for TBK1 and transcription factors activation(Tanaka and Chen, 2012; Zhang et al., 2019). While the tertiary structure and overall architecture of the CBD and TM are well-conserved, the CTT presents significant differences across species. For instance, invertebrate STING lacks a CTT(Wu et al., 2014), with few possible exceptions such as the oyster *Crassostrea gigas* (mollusk) and shrimp *Litopenaeus vannamei* (crustacean)(Li et al., 2024), while the CTT of *Danio rerio* (zebrafish) STING is longer than that of mammals (eg, *Mus musculus* or *Homo sapiens*). In addition, amphibian STING (both Anura and Urodela) exhibit a vestigial CTT, suggestive of secondary loss(de Oliveira Mann et al., 2019; Margolis et al., 2017; Wu et al., 2014). In contrast, despite low amino acid sequence similarity, the CBD of *Crassostrea gigas* (mollusk), *Drosophila melanogaster* (insect), and *Monosiga brevicollis* (choanoflagellate) can all bind CDNs, even though the specific type of CDN differs among species(Cai et al., 2023; Morehouse et al., 2020; Woznica et al., 2021). Interestingly, while most STING homologs present TM, that of *Crassostrea gigas* instead presents a Toll/IL1 receptor (TIR) domain(Morehouse et al., 2020) and a short CTT-like domain of very low sequence similarity with vertebrate CTT. In line with the poor conservation of the CTT, the *bona fide* IFN system originated much later in evolution than the CDN-STING interaction, is restricted to jawed vertebrates, and absent from agnathans and basal chordates(Kasamatsu, 2013). In contrast, NF-κB, as well as 2’3’-cGAMP-induced non-canonical autophagy pathways, are highly conserved in basal metazoans such as *Nematostella vectensis* (cnidarian)*(Margolis et al., 2021)*. Overall, the prevailing hypothesis is thus that STING fulfilled ancestral immune-related functions, *via* autophagy and NF-κB induction, prior to emergence of the IFN system.

STING-mediated lipid metabolism regulation is the main characterized homeostatic function of STING, but its evolutionary conservation has not yet been assessed. In *Mus musculus*, STING controls metabolic homeostasis by inhibiting the enzymatic activity of the fatty acid desaturase 2 (FADS2)(Vila et al., 2022). FADS2 is a Δ-6 desaturase which catalyzes the rate-limiting step in the omega-6 and omega-3 polyunsaturated fatty acid (PUFA) biosynthesis pathway(Sprecher et al., 1995). The action of FADS2 leads to the generation of long chain PUFAs (LC-PUFAs) that serve as ligands of metabolic transcription factors(Grygiel-Gorniak, 2014; Kliewer et al., 1997), or are further processed into signaling lipids called oxylipins(Balvers et al., 2012; Gabbs et al., 2015; Schmitz and Ecker, 2008). *Mus musculus* STING ablation consequently leads to accumulation of FADS2 products, including LC-PUFAs and oxylipins, promoting metabolic rewiring *in vivo(Vila et al., 2022)*. The ability of STING to regulate lipid homeostasis is also documented in human(Mansouri et al., 2022) and in *Drosophila melanogaster(Akhmetova et al., 2021)*. Interestingly, STING haplotypes have been reported in human populations that are deficient for IFN response induction, but competent for lipid metabolism regulation and FADS2 interaction(Mansouri et al., 2022; Vila et al., 2022), suggesting conservation of this homeostatic function. Importantly, this role of STING in the regulation of lipid metabolism is profoundly disturbed in conditions of STING activation(Vila et al., 2022), where STING is displaced from the ER where it regulates FADS2, to the Golgi apparatus and subsequently degraded(Abe and Barber, 2014).

Since STING predates the evolution of IFN-based immunity(Morehouse et al., 2020; Woznica et al., 2021), is present in basal metazoans(Margolis et al., 2021), has been shown to have a critical role in lipid metabolism(Akhmetova et al., 2021; Mansouri et al., 2022; Vila et al., 2022), and in several instances has lost its ability to induce IFN-response secondarily(de Oliveira Mann et al., 2019; Mansouri et al., 2022), we hypothesized that the role of STING in metabolic regulation may be its primordial function. To test this hypothesis, we analyzed the conservation of the FADS2-STING interaction and of the associated functional output.

## RESULTS

### Rationale for the choice of STING homologs

Based on the presence of key domains and on the ability to activate downstream effectors, human STING homologs in eukaryotes can be divided into 3 broad classes. First, the STING homologs found in the close unicellular relatives of metazoans such as choanoflagellates, which can induce autophagy upon activation(Woznica et al., 2021), and for which we chose *Monosiga brevicollis* (*M. brevicollis*) as a representative (Figure 1A). Second, the STING homologs found in basal metazoan phyla, including species such as the sea anemone *Nematostella vectensis* (*N. vectensis*), or in protostomians including arthropods like *Drosophila melanogaster* (*D. melanogaster*) and mollusks like *Crassostrea gigas* (*C. gigas*). These STING proteins reportedly induce NF-κB signaling in response to activation(Hua et al., 2018; Li et al., 2022; Margolis et al., 2021; Martin et al., 2018) (Figure 1A). Third, STING from (jawed) vertebrates, such as *Danio rerio* (*D. rerio*), *Mus musculus* (*M. musculus*), or *Homo sapiens* (*H. sapiens*), which can induce type I IFN responses(Biacchesi et al., 2012; de Oliveira Mann et al., 2019; Ishikawa and Barber, 2008; Wu et al., 2014) (Figure 1A). A notable exception within the vertebrate group is the STING from amphibians such as *Xenopus tropicalis* (*X. tropicalis*), which cannot induce an IFN-response due to absence of a functional CTT(de Oliveira Mann et al., 2019).

**Figure 1.**
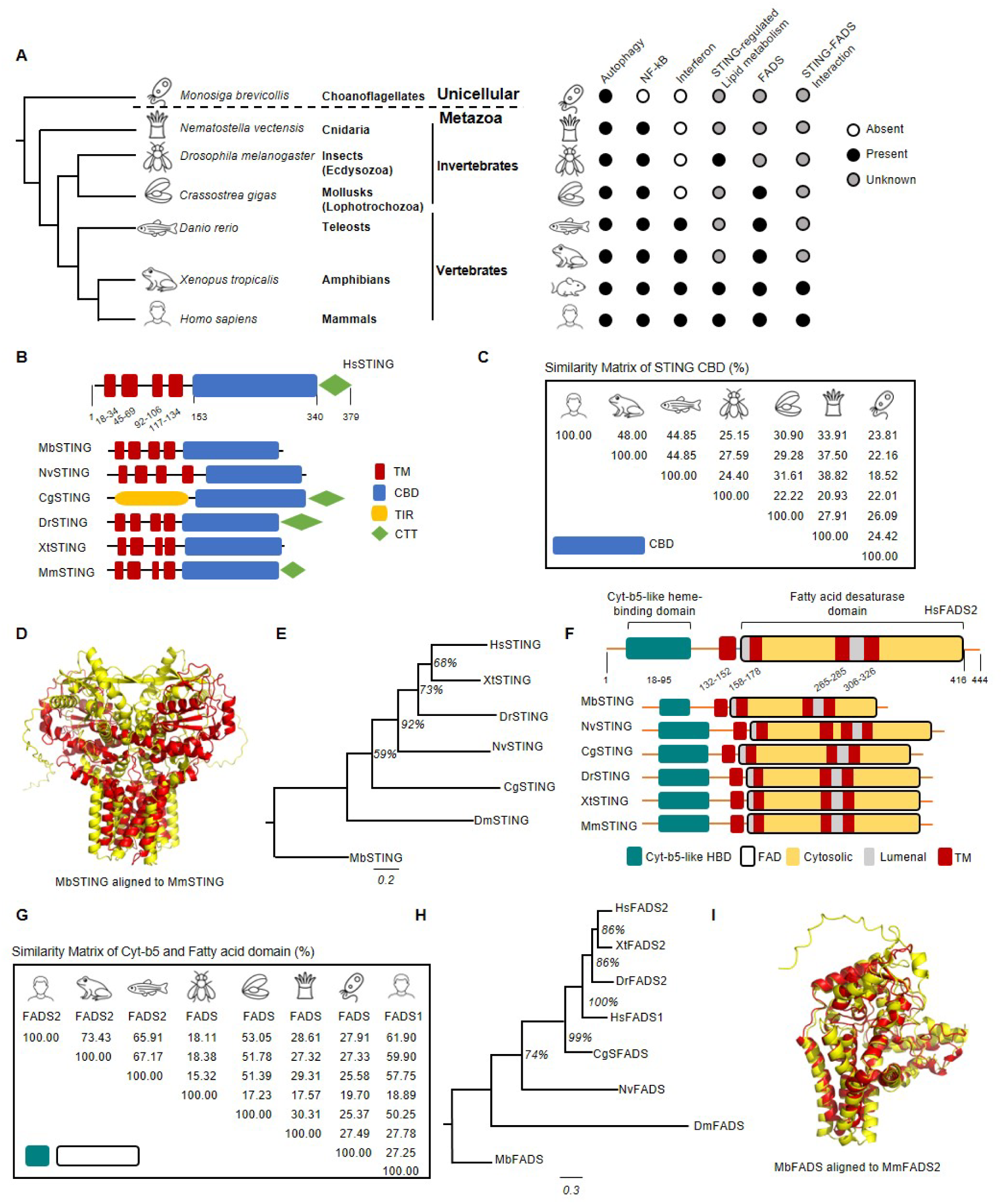
Comparative analyses of STING and FADS homologs. (A) Schematic evolutionary tree of representative species showing phylogenetic relationships (branch lengths are not to scale) with table showing conservation of STING effectors/functions, i.e. STING-dependent autophagy, NF-κB activation, Interferon activation, lipid metabolism regulation and interaction with FADS2 (Orange: present, white: absent, blue: unknown). (B) Structure of STING domains in different lineages. The key to domains representation is shown under the panel. Abbreviations: CTT, C-terminal tail; STING, stimulator of interferon genes; TIR, Toll and IL-1 receptor; CDN, cyclic dinucleotide; CBD: cyclic dinucleotide binding domain. (C) Matrix of sequence similarity of STING CBD domains (from Clustal omega at ebi.ac.uk) (D) Sequence-independent structure-based alignment of MbSTING (red) and MmSTING (yellow) done using Pymol. (E) Evolutionary tree (unrooted) of STING homologs inferred by Maximum Likelihood method and JTT matrix-based model. The tree with the highest log likelihood (-6633.58) is shown. The percentage of trees in which the associated taxa clustered together is shown next to the branches. Evolutionary analyses were conducted in MEGA X. (F) FADS2 and FADS-like proteins domain structure in the representative species. The key to domains representation is shown under the panel. Abbreviations: FAD: fatty acid desaturase; HBD: heme binding domain, TM: transmembrane domain. (G) Matrix of sequence identity of the region comprising CytB5 and FAD domains (from Clustal omega at ebi.ac.uk). (H) Evolutionary tree inferred by Maximum Likelihood method and JTT matrix-based model. The tree with the highest log likelihood (-6652.47) is shown. The percentage of trees in which the associated taxa clustered together is shown next to the branches. Evolutionary analyses were conducted in MEGA X. (I) Structure-based alignment of MbFADS (red) and MmFADS2 (yellow) done using Pymol. Related to supplementary Figure 1 and supplemental Items 1 and 2.

When the domain organization of STING from these species is considered, it is notable that all sequences comprise four N-terminal TMs, with the exception of STING from *D. melanogaster* (DmSTING) which comprises 2-3 predicted TMs(Goto et al., 2018; Wu et al., 2014) and STING from *C. gigas* (CgSTING) which is devoid of TM but rather harbors a TIR domain(Morehouse et al., 2020) (Figure 1B), suggesting that this protein may not localize to the ER. Finally, the CTT is generally absent from STING proteins of invertebrates, and presents the highest variability among vertebrates (Figure 1B). CgSTING however presents a short putative CTT, highly divergent from vertebrate CTT (Figure 1B and Supplementary Item S1).

Excluding the CTT, the STING C-terminal domain – which contains the CDN-binding region (CBD) – exhibits low overall amino acid sequence conservation (Figure 1C), despite the CDN binding motif/pocket itself being highly conserved (Supplementary Item S1). Prediction of the three-dimensional conformation of STING homologs across holozoan evolution shows a similar global architecture (Supplementary Figure S1A). However, structural alignment of the most evolutionary distant STING homologs, namely the choanoflagellate MbSTING and the mouse MmSTING, showed poor overlap of the CBD (Figure 1D). Unrooted evolutionary tree of STING-domains do not mirror the tree of life, consistent with varying pace of domain acquisition, modification, or loss during metazoan evolution (Figure 1E).

We thus selected representative STING genes from each of the three aforementioned groups to assess the conservation of STING-FADS2 interaction, and the role STING in lipid metabolism regulation. Specifically, we used MbSTING for unicellular eukaryotes; NvSTING for basal metazoans; and MmSTING, XtSTING, and DrSTING for vertebrates.

### Identification of FADS2 homologs

FADS2 bears a Δ-6 desaturase activity, which is essential for the desaturation of PUFAs to LC-PUFAs. Δ-6 desaturase activities, essential for lipid metabolism have ancient origins(Castro et al., 2012; Monroig et al., 2022). The general structure of the prototypical FADS2 comprises 3 domains (Figure 1F): an N-terminal cytochrome b5-like heme/steroid binding domain, which acts as the electron donor during desaturation of PUFAs, facilitating the transfer of electrons to the catalytic site; four TM domains that ensure anchoring of FADS2 to the ER and proper positioning of the catalytic site; the C-terminal fatty acid desaturase domain comprising histidine-rich motifs which form the catalytic site for desaturase activity(Blahova et al., 2020).

Putative orthologs of mouse FADS2 have been identified in vertebrates *H. sapiens*, *D. rerio*, *X. tropicalis(Castro et al., 2012)*, but not in *N. vectensis,* or *M. brevicollis*. Through analyses of the genome of these 2 latter species, we identified FADS genes (Figure 1F-G) corresponding to distant homologs of vertebrate FADS1 and FADS2 (Figure 1G-1H). However, the global domain structure is similar in all these proteins, and the cytochrome b5-like domain is very well conserved (Supplementary Item S2). We thus modeled the 3D structure of FADS2 and FADS-like Δ-6 desaturase enzymes to assess the conservation of the protein architecture and ascertain that the identified homologs in *N. vectensis* and *M. brevicollis* are *bona fide* FADS candidates with putative similar functions. Overlay of the models showed that the overall architecture is conserved (Figure 1I and Supplementary Figure S1B).

Taken together, these analyses identify FADS2-related proteins (FADS) in all representative species from the phyla in which we aimed to characterize the STING-FADS interaction.

### STING from multiple species interact with their conspecific Δ6-desaturase enzymes

To test if STING from multiple species can interact with their identified conspecific FADS-like Δ6-desaturase enzymes, codon-optimized STING and FADS2 or FADS-like sequences were hemagglutinin (HA)- and FLAG-tagged respectively (Figure 2A), and inserted into retroviral vectors suitable for expression in mammalian cells. The FADS2 homologs cloned and tested for expression were from *M. brevicollis* (MbFADS), *N. vectensis* (NvFADS), *D. rerio* (DrFADS2), *X. tropicalis* (XtFADS2), and *M. musculus* (MmFADS2) (Figure 2A).

**Figure 2.**
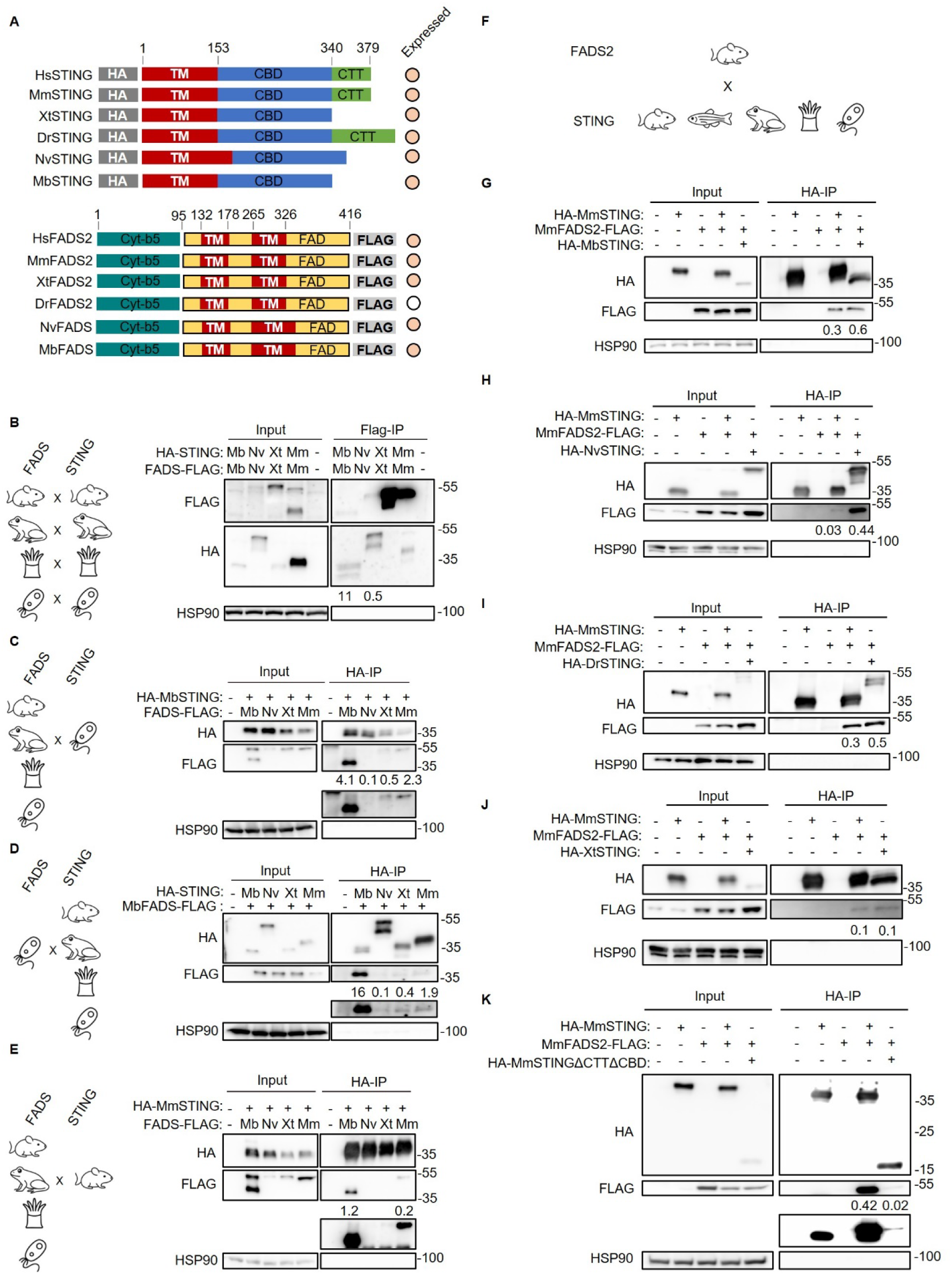
The interaction of STING with FADS is conserved. (A) Top panel: Schematics of HA-tagged STING constructs roughly aligned to HsSTING and their expression status in HEK 293t cells. Bottom panel: Schematics of FLAG-tagged FADS constructs roughly aligned to HsFADS2 and their expression status in HEK 293t cells indicated (orange: present, white: absent). (B) Flag-immunoprecipitation was performed on whole-cell extracts (WCE) from HEK 293T cells transiently co-expressing HA-STING and FADS-FLAG of the indicated species or not. Inputs and eluates from the immunoprecipitation were analyzed by western blot (WB) using indicated antibodies. (C) HA-immunoprecipitation was performed on WCEs from HEK 293T cells transiently co-expressing HA-MbSTING with FADS-FLAG from the indicated species or not. Inputs and eluates from the immunoprecipitation were analyzed as in B. (D) As in C, except that cells co-expressed MbFADS-FLAG with HA-Sting from the indicated species or not. (E) As in C, except that cells co-expressed HA-MmSTING with FADS-FLAG from the indicated species or not. (F) Interaction studies of STING proteins from the indicated species with FADS2 from mouse rendered using clipart. (G) HA-immunoprecipitation was performed on WCEs from HEK 293T cells transiently co-expressing MmFADS2-Flag or not with HA-MbSTING, HA-MmSTING (positive control), or none. Inputs and eluates from the immunoprecipitation were analyzed by WB using indicated antibodies. (H) As in H, except that cells expressed HA-NvSTING. (I) As in H, except that cells expressed HA-DrSTING. (J) As in H, except that cells expressed HA-XtSTING. (K) HA-immunoprecipitation was performed on WCEs from HEK 293T cells transiently co-expressing MmFADS2-FLAG or not with HA-MmSTING (positive control), HA-MmSTINGΔCBDΔCTT, or none. Inputs and eluates from the immunoprecipitation were analyzed by WB using the indicated antibodies. WBs are representative of 3 independent experiments. The normalized binding index (indicated below the co-immunoprecipitated protein) shown for the immunoprecipitation experiments in transient expression cell lines was calculated by normalizing the ratio between band intensities of the immunoprecipitated prey and bait to that of the prey input. Related to supplementary Figure 2

We then tested the ability of MbSTING, NvSTING, and XtSTING to interact with conspecific FADS2 homologs. To this aim, HA-STING and FADS-FLAG constructs were co-expressed in HEK 293T cells prior to FLAG immunoprecipitation (IP), release of bound material by peptide elution, and analysis by WB. MmSTING and MmFADS2 were used as positive controls. Despite low expression levels, we found that MbFADS2 and NvFADS2 co-immunoprecipitated with MbSTING and NvSTING, respectively (Figure 2B), with an enrichment higher than that of MmSTING with MmFADS2 (Figure 2B). These observations indicated that an interaction between STING and a FADS-like protein arose prior to the emergence of metazoans, long before the duplication event that produced FADS1 and FADS2. In contrast, XtSTING did not co-immunoprecipitate with the expressed isoform of XtFADS2, suggesting that either the ability to interact with FADS2 was lost in *X. tropicalis* or that this ability is not harbored by this specific isoform.

We next assessed the ability of the STING and FADS2 homologs to interact with their putative counterparts from distant species. First, HA-STING from *M. brevicollis* was co-expressed with MbFADS, NvFADS, XtFADS2, or MmFADS2 prior to HA-IP and WB analyses. Co-immunoprecipitation of MbFADS, XtFADS2, and MmFADS2 was detected with MbSTING (Figure 2C). No interaction of NvFADS was detected with MbSTING possibly due to very low expression levels (Figure 2C). We next co-expressed MmSTING, XtSTING, NvSTING, and MbSTING with MbFADS prior to HA immunoprecipitation and WB analysis of FADS2 co-immunoprecipitation. We found that MbFADS co-immunoprecipitated with XtSTING and MmSTING and to a lesser extent with NvSTING (Figure 2D). Next, when MmSTING was co-expressed with MbFADS, NvFADS, and XtFADS2, we found strong co-immunoprecipitation of MbFADS with MmSTING, while weak interaction of XtFADS2 was detected (Figure 2E). Finally, we assessed whether STING from different species are capable of interacting with MmFADS2 (Figure 2F). Co-expressing MbSTING, NvSTING, DrSTING, and XtSTING with MmFADS2 prior to HA-immunoprecipitation and WB analysis of FADS2 interaction showed that all tested homologs bear the capacity to interact with MmFADS2 (Figure 2G-J).

Together, these data suggest that STING from multiple species bear the ability to interact with FADS-related proteins from distant species, although the strength of the detected interaction is variable. It is notable that the interaction of the FADS from a non-metazoan, *i.e.* MbFADS, was strongest with the mammalian MmSTING, in line with the idea that the ability to interact with FADS-like Δ6-desaturase enzymes is a conserved ancient function of STING , although the strength of the interactions did not correlate very well with evolutionary proximity.

### The CTT is dispensable and TM domain sufficient for interaction of STING with FADS2

Our analyses show that STING from basal organisms lacking a CTT are able to interact with FADS2, suggesting that the CTT is likely dispensable for this interaction. Previous work(Vila et al., 2022) has shown that the TM domain of STING is essential for interaction with FADS2. To further investigate which domains are required for interaction between FADS2 and STING, we generated HA-tagged versions of STING homologs with unusual domain architecture, namely from the bacterium *Flavobacteriaceae sp.* (FsSTING, *cap13*) and mollusk *C. gigas* (CgSTING). In addition, we generated MmSTING and DrSTING mutants where the CTT is ablated (MmSTINGΔCTT and DrSTINGΔCTT respectively), as well as a MmSTING mutant comprising the TM domain alone (MmSTINGΔCTTΔCBD) (Supplementary Figure S2A). The interaction of these alleles with MmFADS2 was subsequently tested in co-immunoprecipitation experiments.

FsSTING (Supplementary Figure S2B) and CgSTING (Supplementary Figure S2C) were unable to interact with FADS2, confirming that ER anchoring is likely essential for interaction of STING with FADS2(Vila et al., 2022). Indeed, FsSTING does have two TM domains, but owing to the absence of ER in bacteria, it can be speculated that the FsSTING localizes to other compartments, while CgSTING lacks TM domains altogether. In contrast, we found that ablation of the CTT led to enhanced interaction of MmSTINGΔCTT and DrSTINGΔCTT with MmFADS2, as compared to the WT alleles (Supplementary Figure S2D-E). Finally, we found that MmSTINGΔCTTΔCBD retained the ability to interact with MmFADS2 (Figure 2K), and so did the STING homolog of *Drosophila melanogaster* (DmSTING) whose CBD has very low sequence similarity with other STING homologs (Supplementary Figure S2F).

Together these data show that the localization of STING at the ER membrane is essential for the interaction with FADS2, and that the TM region is both essential and sufficient for this interaction, while the CTT is dispensable. This further supports that the interaction of FADS2 and STING predated the emergence of the CTT in the STING of jawed vertebrates.

### STING homologs interact with Mus musculus FADS2 and are functional in murine cells

In order to evaluate the capacity of the STING homologs from different taxonomic groups to regulate FADS2 activity, we next engineered mouse embryonic fibroblasts (MEF) knockout for Sting (MEF*^Sting-/-^*) stably expressing STING homologs from representative species *M. brevicollis*, *N. vectensis*, *D. rerio,* and *X. tropicalis*: MEF*^MbSTING^*, MEF*^NvSTING^*, MEF*^DrSTING^*, MEF*^XtSTING^*, and MEF*^MmSTING^*, respectively. Immunoprecipitation of HA-tagged STING homologs was conducted coupled to analysis of endogenous FADS2 co-immunoprecipitation, showing that all tested homologs interacted with endogenous MmFADS2 (Figure 3A). This not only comforts that the interaction between murine FADS2 and STING homologs is well-conserved, but also suggests that the constructed heterologous cell lines are suitable for assessment of STING function.

**Figure 3.**
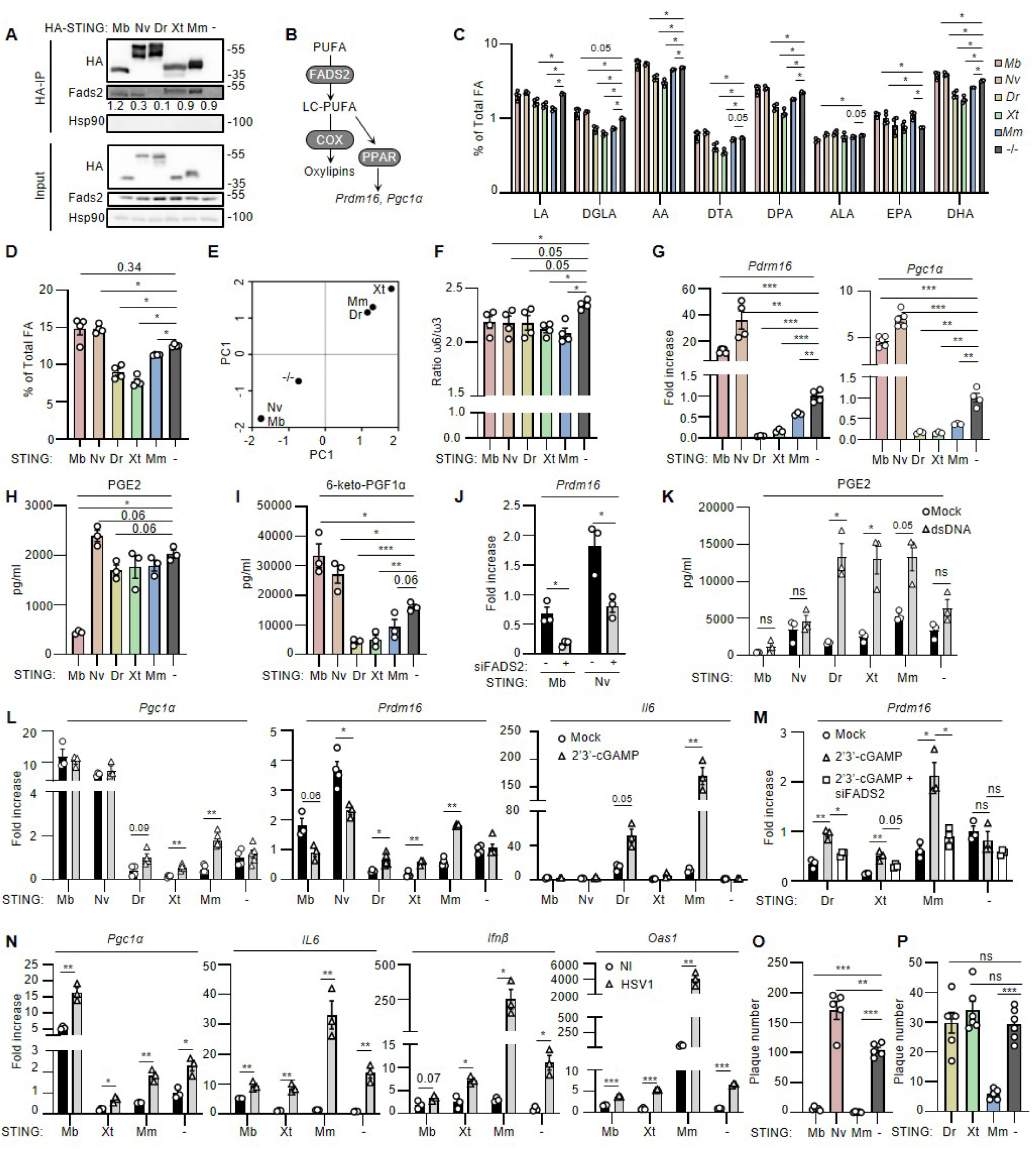
STING homologs differentially modulate PUFA pools. (A) HA-immunoprecipitation was performed on WCEs from MEF*^Sting-/-^* cells stably expressing HA-tagged STING homologs from the indicated species or not. Input and immunoprecipitated material were analyzed by WB using the indicated antibodies. (B) Scheme indicating the products downstream of FADS2 activity. In brief, FADS2 generates LC-PUFAs from PUFA substrates. LC-PUFAs can operate as ligands of transcription factors ultimately leading to transcription of *Prdm16* an *Pgc1α*, or be metabolized by enzymes such as cyclooxygenases (COX) to give rise to oxylipins. (C) Relative quantities of the indicated PUFAs and LC-PUFAs in MEF*^Sting-/-^* cells expressing HA-STING homologs from the indicated species measured using LC-MS and expressed as percentage of total fatty acids. Graph presents the mean (±standard error of the mean (SEM)) of n=4. (D) Summation of the LC-PUFA products of FADS2 expressed as percentage of total fatty acids from the LC-PUFAs measured in C. Graph presents the mean (±SEM) of n=4. (E) PC score plot from principal component analysis performed using relative quantities of FADS2 products as variables using the data from C. (F) Ratio of total omega-6 to omega-3 PUFAs measured in C. Graph presents the mean (±SEM) of n=4. (G) RT-qPCR analysis of *Prdm16* and *Pgc1α* mRNA levels in the cell lines used in C, normalized to *Hsp90* and expressed as fold increase over MEF*^Sting-/-^.* Graph presents the mean (±SEM) of n=3-4. (H-I) PGE2 and 6-keto-PGF1α concentrations in the cell lines used in C, were quantified by ELISA and normalized to total protein concentration in indicated cell lines. Graph presents the mean (±SEM) of n=3. (J) *Prdm16* mRNA levels 48 hrs after transfection of MEF^Sting-/-^ cells expressing HA-STING homologs from the indicated species with 10 nM siFADS2 or control, normalized to *Hsp90* and expressed as fold increase over mock transfected. Graph presents the mean (±SEM) of n=3. (K) PGE2 levels were measured by ELISA in cells lines treated with 1 µg/ml of 80nt dsDNA for 24 hrs. Graph presents the mean (±SEM) of n=3. (L) RT-qPCR analysis of *Pgc1α, Prdm16,* and *Il6* mRNA levels in cell lines expressing HA-tagged STING homologs from the indicated species, stimulated or not with 14 µM 2’3’-cGAMP for 9h, normalized to *Hsp90* and expressed as fold increase over MEF*^Sting-/-^* unstimulated. Graph presents the mean (±SEM) of n=3-5. (M) *Prdm16* mRNA levels in MEF^Sting-/-^ cells expressing HA-STING homologs from the indicated species transfected with 10 nM siFADS2 or control followed by stimulating or not with 14 µM 2’3’-cGAMP for 9h, normalized to *Hsp90* and expressed as fold increase over MEF*^Sting-/-^* siControl unstimulated. Graph presents the mean (±SEM) of n=3. (N) Cell lines expressing MbSTING, XtSTING, and MmSTING were infected with HSV-1 for 6 and 16 hours prior to gene expression analysis. Graph presents the mean (±SEM) of n=3. (O) Cell lines expressing indicated STING homologs were infected with HSV-1 prior to plaque analysis at 48 hours post infection. Graph presents the mean (±SEM) of n=5. (P) As in O, except that cell lines expressing vertebrate STING homologs were used. Graph presents the mean (±SEM) of n=5-6. WBs are representative of 3 independent experiments. All graphs are means ± SEM from at least 3 independent experiments. p-values were determined either by Student’s t test or by Mann-Whitney test. *p < 0.05, **p < 0.01, ***p < 0.001. Related to supplementary Figure 3-4

We next tested the functionality of these heterologous murine cells expressing STING from distant species, by assessing their response to stimulation with 2’3’-cGAMP, followed by analysis of subcellular localization by immunofluorescence, oligomerization by native electrophoresis, and pathway activation by immunoblotting. Immunofluorescence analyses showed that all the homologs are localized in the ER membrane in the absence of stimulation, as suggested by colocalization with calnexin (Supplementary Figure S3A-B). Stimulation with 2’3’-cGAMP promoted their displacement from the ER, as suggested by decreased co-localization with calnexin (Supplementary Figure S3A-B). Of note, the subcellular localization of endogenous MmFADS2 was also assessed, showing no change in localization following treatment with 2’3’-cGAMP (Supplementary Figure S3A). When oligomerization in response to 2’3’-cGAMP was assessed, increase in tetramers and higher order oligomers were observed for NvSTING (Supplementary Figure S3C), DrSTING, and MmSTING (Supplementary Figure S3D), after 1-3 hours of treatment with 2’3’-cGAMP. For XtSTING, dimer and tetramers were not clearly detectable, but an increase in oligomers was observed at 1-3 hours post treatment (Supplementary Figure S3D). In contrast, MbSTING showed an increase in tetramers 1-3 hours post treatment, but no clear increase in higher order oligomers (Supplementary Figure S3C). Finally, the induction of autophagy, the most ancient and well-conserved pathway downstream of STING activation(Gui et al., 2019; Woznica et al., 2021), was also detected showing that all homologs induced autophagy to levels similar to that observed in cells expressing MmSTING (Supplementary Figure S3E-F).

Together these data show that the STING homologs expressed in MEF respond to stimulation with 2’3’-cGAMP, indicating that they are able to perform their conserved innate immunity-related functions, such as inflammatory response and autophagy induction.

### STING homologs regulate lipid metabolism differently across species

We next assessed whether STING homologs from distant species, which are capable of interacting with MmFADS2, also regulate its activity, focusing on MbSTING, NvSTING, DrSTING, and XtSTING. To this aim, we performed fatty acid profiling, analyzed transcriptional activation of metabolic genes downstream of LC-PUFA accumulation, and measured oxylipins in the heterologous cell systems that we generated (Figure 3B).

We quantified the detectable PUFAs and LC-PUFAs by liquid chromatography coupled to mass spectrometry (LC-MS) (Figure 3C). This showed that levels of all the detected omega-6 PUFAs *i.e.* linoleic acid (LA), dihomo-γ-linolenic acid (DGLA), arachidonic acid (AA), docosatetraenoic acid (DTA), and docosapentaenoic acid (DPA) were significantly reduced in cells expressing vertebrate STING homologs (DrSTING, XtSTING and MmSTING), while among the omega-3 PUFAs, only docosahexaenoic acid (DHA) showed a significant decrease (Figure 3C). In contrast, invertebrate STING (MbSTING and NvSTING) expression led to increased levels of DGLA, AA, DTA, DPA, eicosapentaenoic acid (EPA), and DHA (Figure 3C).

The summation of the products of FADS2 detectable in our assay, reflecting its activity, showed that the expression of MmSTING decreased FADS2 activity as previously reported(Vila et al., 2022) (Figure 3D). Similarly, DrSTING and XtSTING also decreased the levels of FADS2 products (Figure 3D). On the contrary, the levels of FADS2 products measured in cells expressing MbSTING and NvSTING not only increased as compared to cells expressing MmSTING, but also compared to *Sting-/-* cells (Figure 3D). This distinct behavior of vertebrate and invertebrate STING homologs is illustrated by principal component analysis (PCA) with the relative quantities of PUFAs as variables (Figure 3E). Cells expressing STING from representative species classified into two distinct clusters both markedly different from the *Sting-/-* cells: STING from the invertebrate or unicellular eukaryote species have an activating effect while vertebrate STING have an inhibiting effect on FADS2 activity (Figure 3E). Regardless of this, the ratio of total omega-6 to omega-3 PUFAs was reduced for all tested homologs (Figure 3F), potentially indicating that they skew FADS2 activity towards enhanced processing of omega-6 PUFAs.

PUFAs strongly activate peroxisome proliferator-activated receptors (PPAR) transcription factors involved in regulation of glucose and lipid metabolism(Kliewer et al., 1997), upregulating the expression of thermogenic regulators like PPARγ coactivator 1-alpha (*Pgc1α*) and PR domain-containing 16 (*Prdm16*)(Seale et al., 2007). Our previous work(Vila et al., 2022) has shown that one of the effects of STING ablation in mice is the upregulation of these genes due to accumulation of LC-PUFAs resulting from increased FADS2 activity. Therefore, we used their messenger ribonucleic acid (mRNA) levels in the heterologous cell lines as a read-out for MmFADS2 inhibition by the respective heterospecific STING (Figure 3B). We found that all vertebrate STING homologs efficiently downregulated *Prdm16* and *Pgc1α* expression (Figure 3G). Surprisingly, mRNA levels of both *Prdm16* and *Pgc1α* were higher than MEF*^Sting-/-^*for the two other STING homologs (Figure 3G). Finally, we measured oxylipins that are metabolites downstream of AA generated by FADS2 activity (Figure 3B), namely prostaglandin E2 (PGE2) and 6-keto-PGF1, which is a stable isoform of PGI2. We found that the expression of vertebrate STING led to a decrease of PGE2 and 6-keto-PGF1 (Figure 3H-I). Expression of other STING homologs caused an increase of both oxylipins in the case of NvSTING and an increase of 6-keto-PGF1 and decrease of PGE2 for MbSTING (Figure 3H-I). Since increase of FADS2 activity by STING is unheard of, we verified that the increased activation of Prdm16 activity is dependent on FADS2 by using small interfering RNAs (siRNAs) targeting *Fads2* (Supplementary Figure S4A). We found that silencing FADS2 led to decreased expression of *Prdm16* in MEF*^MbSting-/-^* and MEF*^NvSting-/-^* (Figure 3J and Supplementary Figure S4B). Altogether, these data indicate that while vertebrate STING homologs are able to decrease the levels of FADS2 products, homologs from invertebrate or unicellular eukaryote species promote an increase of FADS2-derived metabolites in general.

Our interaction analyses showed that the CTT was dispensable for interaction of STING with FADS2. We assessed if the presence of a CTT would alter FADS2 activity. To this aim, MEF*^Sting-/-^* engineered to stably express MmSTING, DrSTING, MmSTINGΔCTT, and DrSTINGΔCTT were subjected to fatty acid profiling (Supplementary Figure S4C). The levels of omega-6 PUFAs AA, DTA, and DPA were significantly reduced upon ablation of the CTT. The omega-3 PUFAs EPA and DHA also showed reduction in the ΔCTT mutants. Summation of FADS2 products showed that ablation of the CTT causes a slightly higher decrease of FADS2 products (Supplementary Figure S4D). PCA (Supplementary Figure S4E) and calculation of the omega-6 to omega-3 PUFA ratio (Supplementary Figure S4F) showed no significant variation. Finally, gene expression analysis showed a downregulation of *Prdm16* when MmSTINGΔCTT is expressed as compared to MmSTING, but not when the CTT of DrSTING is ablated (Supplementary Figure S4G). No significant change in *Pgc1α* mRNA levels was observed upon ablation of the CTT of either species (Supplementary Figure S4G). This suggests that the CTT is not just dispensable for the interaction with FADS2, but probably also impedes the inhibitory effect of STING to some extent, in particular in the case of MmSTING.

### STING activation disrupts lipid metabolism regulation differently across species

Previous work indicated that STING activation by agonists disrupts the inhibition of FADS2 activity in murine cells, notably through the displacement of STING from the ER compartment(Vila et al., 2022). We questioned whether a similar release of STING-mediated inhibition of FADS2 activity can be witnessed for STING homologs after stimulation.

We assessed the changes in STING homolog-mediated FADS2 activity modulation upon stimulation with dsDNA which stimulates production of 2’3’-cGAMP by cGAS. Measurement of PGE2 in this context showed no change upon dsDNA stimulation in cells expressing NvSTING and MbSTING, while an increase was observed for cells expressing vertebrate STING homologs (Figure 3K). Next, cells were challenged with 2’3’-cGAMP followed by analysis of the expression of *Prdm16* and *Pgc1α* as a surrogate for FADS2 activation. *Il6* expression was monitored to assess the pro-inflammatory status of cells. We found that 2’3’-cGAMP induced *Il6* expression in cells expressing DrSTING and MmSTING (Figure 3L), consistent with their ability to trigger NF-kB activity (Supplementary Figure S3F). In the cells expressing STING from vertebrate classes, 2’3’-cGAMP stimulation led to a significant increase in the expression of *Prdm16* and *Pgc1α,* suggesting alleviation of their inhibition of FADS2 activity (Figure 3L). In contrast, no significant change to *Pgc1α* levels was observed for NvSTING and MbSTING expressing cells, while a reduction of *Prdm16* and was observed for both homologs (Figure 3L). We next verified whether the increased expression of metabolic response genes was dependent on FADS2 by using siRNAs to knockdown FADS2 prior to stimulation by 2’3’-cGAMP. We found that the increase in Prdm16 levels was reverted to non-stimulated levels upon knockdown (Figure 3M).

We next questioned whether infection by a dsDNA genome-harboring virus, such as the human herpes simplex virus 1 (HSV-1), known to promote cGAS-dependent 2’3’-cGAMP production, can induce similar alterations of FADS2 activity in the presence of STING homologs. To this aim, we used cells expressing MbSTING, as representative of non-vertebrate models, and cells expressing XtSTING and MmSTING as representative of vertebrates. This choice of STING homologs also facilitated the isolation of IFN-independent processes. Gene expression analysis showed that MbSTING expression led to an increase of *Pgc1α*, *Il6* and of the *Oas1* IFN response gene, and a tendency for increase of *Ifnβ* at 16 hours post infection. Similarly, an increase of *Pgc1α*, *Il6*, *Ifnβ,* and *Oas1* was observed for XtSTING expressing cells. For both homologs, the observed increase in inflammation-related gene expression (*Il6*, *Ifnβ,* and *Oasl1*) was not significantly different from what is observed in STING-deficient cells and below the response induced in MmSTING-expressing cells (Figure 3N). This data suggests that activation of the tested STING homologs by endogenously produced 2’3’-cGAMP can induce a weak, but reproducible increase of metabolic gene activation alongside the prototypical STING associated inflammatory and type I IFN response.

Finally, we interrogated whether the STING homologs tested in this study may be able to promote the establishment of an antiviral status, despite low level of type I IFN response induction, through their ability to modulate lipid metabolism. To this aim, we infected cells expressing MbSTING or NvSTING with HSV-1 and measured viral replication through plaque assays. We found that, expression of MbSTING, but not of NvSTING, restricted viral replication (Figure 3O). Next, cells expressing vertebrate STING homologs were infected with HSV-1 prior to measurement of viral plaques. We found that both DrSTING and XtSTING failed to restrict HSV-1 plaque formation to the contrary of MmSTING (Figure 3P). Thus, the ability of DrSTING and XtSTING to inhibit FADS2 activity does not correlate with an ability to restrict HSV-1 infection. These data further support that the ability to regulate lipid metabolism does not correlate with restriction of DNA virus replication.

Altogether, our data show that although the ability of STING to regulate lipid metabolism is highly conserved, it does not appear to bear an ancient anti-viral function through inflammatory lipid-derived mediator regulation.

### Human STING haplotypes regulate FADS2 activity

To investigate whether different human STING haplotypes bind FADS2, we generated codon-optimized HsFADS2-FLAG sequence and HA-HsSTING sequences for STING-R232H, STING-AQ (G230A, R293Q), and STING-HAQ (R71H-G230A-R293Q) haplotypes, along with a STING-R232H-ΔCTT mutant (Δ342-379) (Figure 4A). These sequences were co-expressed into HEK 293T cells, prior to HA-immunoprecipitation. The bound material released by peptide elution was then analyzed by WB (Figure 4B). Interestingly, we observed that co-expression of FADS2 with HA-HsSTING-R232H led to a stabilization of the total FADS2 protein levels in the input material (Figure 4B). We calculated the ratio of FADS2 to STING in the eluate, followed by normalization to FADS2 levels in the input to account for variance in the level of FADS2 and STING haplotype/mutant expression. Although all of the tested haplotypes/mutant of HsSTING bound HsFADS2, the HAQ haplotype showed comparatively weaker interaction (Figure 4B). We also confirmed that like in other vertebrates, the CTT of HsSTING is dispensable for FADS2 binding (Figure 4B).

**Figure 4.**
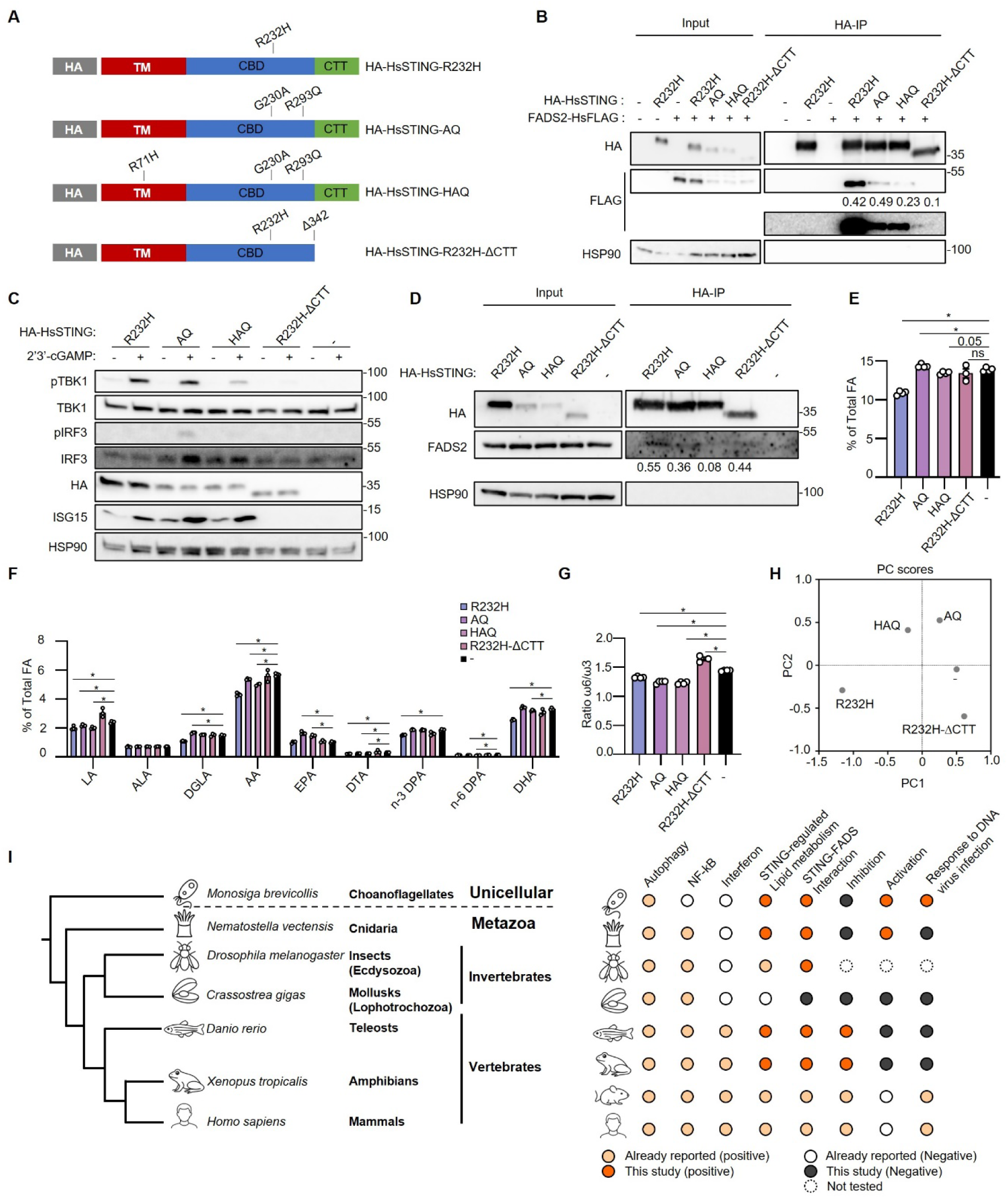
Human STING haplotypes regulate PUFA pools. (A) Schematics of HA-tagged constructs of HsSTING haplotypes and mutants. (B) HA-immunoprecipitation was performed on whole-cell extracts (WCEs) from HEK 293T cells transiently co-expressing HsFADS2-FLAG with one of the indicated HA-tagged HsSTING haplotypes/mutants or none. Inputs and eluates from the immunoprecipitation were analyzed by western blot (WB) using the indicated antibodies. (C) WCEs of ThP-1*^Sting-/-^* cell lines stably expressing the indicated HA-HsSTING haplotype/mutant stimulated or not with 14 µM 2’3’-cGAMP for 6 hrs were analyzed by WB using the indicated antibodies. (D) HA-immunoprecipitation was performed on WCEs from the same cell lines as in C. Input and immunoprecipitated material were analyzed by WB using the indicated antibodies. (E) Summation of the long-chain polyunsaturated fatty acids (LC-PUFA) products of FADS2 expressed as percentage of total fatty acids measured in the same cell lines as in C using liquid chromatography–mass spectrometry (LC-MS). Graph represents the mean ± SEM of n=3-4. (F) Relative quantities of the indicated PUFAs and LC-PUFAs expressed as percentage of total fatty acids as measured in E. Graph represents the mean of n=3-4. (G) Ratio of total omega-6 to omega-3 PUFAs measured in E. Graph presents the mean (±SEM) of n=3-4. (H) PC score plot from principal component analysis (PCA) performed using the relative quantities of PUFAs as variables using the data from E. (I) Scheme summarizing the data obtained in the study. WBs are representative of 3 independent experiments. All graphs are means ± SEM from at least 3 independent experiments. P-values were determined by Student’s t test. *p < 0.05, **p < 0.01, ***p < 0.001. The normalized binding index (indicated below the co-immunoprecipitated protein) shown for the immunoprecipitation experiments in transient expression cell lines was calculated by normalizing the ratio between band intensities of the immunoprecipitated prey and bait to that of the prey input. The binding index (indicated below the co-immunoprecipitated protein) shown for the immunoprecipitation experiments in stable expression cell lines is the ratio between band intensities of the immunoprecipitated prey and bait.

Next, we checked if the interaction takes place in THP-1 human myeloid cells. THP-1*^Sting-/-^* cells were engineered to stably express the *Hs*STING haplotypes, allowing the generation of THP-1*^HsSTING-R232H^*, THP-1*^HsSTINH-AQ^*, THP-1*^HsSTING-HAQ^*, and THP-1*^HsSTING-R232H-ΔCTT^* cell lines. We first tested the functionality of these alleles in THP-1 cells by performing 2’3’-cGAMP transfection, followed by assessment of the phosphorylation of TBK1 and IRF3 and levels of the ISG15 IFN response gene. We found that all haplotypes efficiently allowed activation of TBK1 (Figure 4C). HsSTING-R232H and HsSTING-HAQ presented a lower ability to induce IRF3 phosphorylation as compared to STING-AQ, although all 3 haplotypes could induce ISG15 upregulation (Figure 4C). HsSTING-R232HΔCTT on the other hand lost its ability to induce activation of downstream effectors (Figure 4C).

Next, we used these cell lines to perform HA-IP and found that all haplotypes interacted with FADS2, with a weaker interaction for the HAQ haplotype (Figure 4D), similar to what was observed in HEK 293T cells (Figure 4B). This differential binding of FADS2 by AQ and HAQ haplotypes possibly arises from the single point mutation R71H, which is known to increase the polar solvent exposure of STING (Mansouri et al., 2022; Morere et al., 2022). Moreover, this residue is part of the highly critical TM2–TM3 linker region of STING, which interacts with fatty acyl–CoA(Mansouri et al., 2022). Finally, we also confirm that the CTT is dispensable for interaction with FADS2 (Figure 4D).

We next performed fatty acid profiling and quantification of mRNA levels of genes downstream of perturbation in fatty acid levels, in order to investigate how the human STING haplotypes regulate fatty acid metabolism. Detectable PUFAs were quantified by LC-MS. The summation of detectable products of FADS2, reflecting its enzymatic activity, showed that cells expressing the R232H haplotype had significantly lower LC-PUFAs, while that of AQ was marginally higher than THP-1*^Sting-/-^* (Figure 4E). Levels of major omega-6 PUFAs LA, DGLA, AA, and DTA, were all significantly reduced in R232H compared to THP-1*^Sting-/-^*, but only n-3 DPA and DHA were found to be reduced among omega-3 PUFAs (Figure 4F). For the AQ and HAQ haplotypes, majority of the omega-6 i.e. LA, AA, DTA, and n-6 DPA were reduced, however omega-3 PUFA levels were either unchanged or higher than THP-1*^Sting-/^* (Figure 4F). The R232H-ΔCTT mutant did not show any significant changes in terms of PUFAs levels (Figure 4E-F). All of the 3 haplotypes were able to significantly reduce the omega-6 to omega-3 PUFAs ratio, while the R232H-ΔCTT mutant slightly increased the omega-6 to omega-3 PUFAs ratio (Figure 4G). PCA also illustrated that the R232H presented a distinct behavior, as compared to other haplotypes (Figure 4H). These data show that human STING haplotypes all bear the capacity to regulate PUFA metabolism, but also highlight that the CTT may play a role in this interaction.

Altogether our results point towards the conservation of the ability of STING to regulate PUFA metabolism throughout evolution, which therefore represents a primordial function of this protein (Figure 4I).

## DISCUSSION

In this study, we conducted a thorough analysis of the evolutionary conservation of the interaction of STING with FADS proteins, showing that the ability of STING to interact with FADS-like enzymes and to regulate lipid metabolism arose early in evolution, before the emergence of metazoans. Since prokaryotes also possess STING homologs(Morehouse et al., 2020) along with multiple FADS-like desaturase enzymes(Alonso et al., 2003; Starikov et al., 2022; Tao et al., 2025), the interaction between the two proteins might also exist in prokaryotic cell membrane in an archaic form. Consistent with this ancient role, the CTT domain of STING, which is restricted to vertebrates, is dispensable for the interaction with FADS2 and the regulation of FADS2 activity. While lipid metabolism is virtually present in all living cells, activities of FADS-like enzymes had not yet been reported in all species. We here identify FADS2 homologs in unicellular eukaryotes that contain domains similar to those present in vertebrate FADS2, and possess the ability to interact with STING. While the direct activity of these enzymes should be measured to assert their role in PUFA processing, our data support previous observations that FADS-domain containing enzymes in basal metazoans and unicellular eukaryotes can use PUFAs as substrates(Degraeve-Guilbault et al., 2020; Monroig et al., 2013; Surm et al., 2018).

Duplication events are believed to have given rise to FADS1 and FADS2 in early vertebrates(Castro et al., 2012). These desaturases present distinct desaturase activities, with FADS1 bearing Δ5 activity while FADS2 bears Δ6 activity, and both enzymes cooperating to generate LC-PUFAs(Alonso et al., 2003). Invertebrates rather mostly rely on multifunctional desaturases that can perform both Δ5 and Δ6 activities(Castro et al., 2012; Pereira et al., 2003). Interestingly, zebrafish has secondarily lost its FADS1 and relies on a multifunctional FADS2 enzyme(Abdul Hamid et al., 2016). These differences in PUFA management between vertebrates and invertebrates may account for differential outputs of STING-dependent regulation of FADS-like enzymes. Indeed, we found that STING homologs regulate lipids differently across species from different lineages. In vertebrates, STING efficiently inhibits the desaturation of PUFAs into LC-PUFAs, likely reflecting decreased FADS2 activity. In contrast, we found that STING from unicellular eukaryotes and basal metazoans promote an enhancement of FADS2 products, and subsequent higher activity of tested metabolic transcription factors. This suggests that these STING homologs regulate FADS2 activity differently, or that they may recruit other metabolic enzymes, ultimately modifying metabolic outputs. In this line, recent proteomic studies revealed a diverse array of enzymes involved in lipid metabolism(Motani and Kosako, 2020) in the proximitome of STING. The interaction of STING with these other lipid metabolic enzymes and its evolution remains to be investigated.

Our study highlights an ancient role of STING as a lipid metabolism regulator, prior to the emergence of the IFN system. This raises questions regarding how and why this protein involved in metabolism has been co-opted by the IFN-based innate immune system, and how this dual function of STING conferred an evolutionary advantage. Importantly, lipids generated by FADS2 activity can be processed into signaling lipids, also called oxylipins, that bear signaling properties and can regulate inflammatory processes(Chistyakov et al., 2020; Parchem et al., 2024). Therefore, by coordinating PUFA processing concertedly with type I IFN responses, as well as other innate immune pathways, STING may allow for better fine-tuning of signaling output. In this line, the prevalence of STING haplotypes deficient for type I IFN signaling in some human populations(Mansouri et al., 2022), without reported immune dysfunction(Jin et al., 2011), suggests that the lipid metabolism-regulating function of STING is sufficient for homeostasis. Deeper assessment of the molecular mechanisms through which oxylipins resulting from the concerted action of FADS2 and other enzymes may affect viral infections is therefore warranted to understand whether the function of STING in lipid metabolism regulation may also serve an antiviral function. Additionally, some studies have suggested that STING-mediated regulation of PUFA metabolism may also contribute to antiviral defense by inhibition of viral replication machinery of positive-stranded RNA viruses that rely on ER-derived membranes. This is achieved by lipid peroxidation (LPO)-mediated disruption of membranes, a process that is highly FADS2-dependent and parallels ferroptosis(Yamane et al., 2022; Yamane et al., 2014).

STING is a major therapeutic target, for which a plethora of agonists has been developed with the aim of boosting inflammatory responses in immunosuppressed contexts such as cancer(Hines et al., 2023; Le Naour et al., 2020) (Wang et al., 2024). Our study sheds light onto the way in which STING and FADS2 interact, the conservation of those interactions, as well as the potential metabolic consequences of interfering with STING activity. This work therefore opens a new perspective for the development of therapeutics that would allow dissociating STING-associated inflammatory pathways from metabolic side-effects, or to the contrary contend with STING-dependent lipid metabolism without causing type I IFN responses activation. This is of particular importance when considering the previously established link between STING and ferroptosis(Chen et al., 2023; Li et al., 2021; Wang et al., 2025), which has been described to rely on FADS2 activity and lipid metabolites(Li et al., 2025; Lorito et al., 2024; Yamane et al., 2022). Investigating this interconnection may provide a better understanding of STING-dependent cell death processes in health and diseases.

### Limitations of the study

Knockout of STING in cognate organisms could help demonstrate the inhibitory role of STING homologs in the regulation of lipid metabolism. In addition, infection assays could be conducted with additional viruses, including RNA viruses.

## Supporting information

Supplementary Items

## RESOURCE AVAILABILITY

### Lead contact

Requests for further information and resources should be directed to and will be fulfilled by the lead contact, Nadine Laguette.

### Materials availability

All unique/stable reagents generated in this study are available from the lead contact with a completed materials transfer agreement.

### Data and code availability

- Original data reported in this paper will be shared by the lead contact upon request.
- This paper does not report original code.
- Any additional information required to reanalyze the data reported in this paper is available from the lead contact upon request.

## ACKNOWLEDGMENTS

We thank all members of the Molecular Basis of Inflammation laboratory for their critical reading of this manuscript. We thank Andrei Turtoi for technical discussion of PUFA handling. We thank Dimitrios Vlachakis for conceptual discussions. We thank Soren Paludan for the gift of MEF and MEF*^Sting-/-^*. We acknowledge the MRI imaging facility, member of the national infrastructure France-BioImaging infrastructure supported by the French National Research Agency (ANR-10-INBS-04, “Investments for the future”).

This work was co-funded by the European Union (ERC, SENTINEL 101087092 to NL). Views and opinions expressed are however those of the author(s) only and do not necessarily reflect those of the European Union or the European Research Council. Neither the European Union nor the granting authority can be held responsible for them. This work was also co-funded by: the ANR [AlphA(NL)]; LA Ligue pour la recherche contre le cancer; the Agence Nationale de Recherche sur le SIDA et les Hépatites virales (ANRS) [ECTZ117448 (NL)], the I-SITE Excellence Program of the University of Montpellier, under the Investissements France 2030 [RETTiNA (NL), ChoiCe (NL)], the Fondation ARC [ARCPJA2021060003720 COPALYS (NL)]. SG benefited from a PhD fellowship from the LabMUSE EpiGenMed and from the Fondation pour la Recherche medicale (FRM, FDT202404018184). YMN benefitted from Posdoctoral funding from the FRM (SPF202409019744). HC was funded by the Fondation ARC and by LA Ligue pour la recherche contre le cancer. This research was funded by the Agence Nationale de la Recherche ANR-21-35–0019CE (LipofishVac, PB) and by institutional grants from INRAE (PB).

## AUTHOR CONTRIBUTIONS

Conceptualization: N.L.; Methodology: S. Guha; Investigation: S. Guha, S. Grégoire, N.A., P.B., I.K.V., Y.M.N., M.C., J.R.; Visualization: S. Guha, P.B., I.K.V., N.L.; Funding acquisition: N.L.; Project administration: N.L.; Supervision: N.L., I.K.V.; Writing – original draft: N.L., S. Guha.; Writing – review & editing: N.L., S. Guha

## DECLARATION OF GENERATIVE AI AND AI-ASSISTED TECHNOLOGIES

During the preparation of this work, the author(s) used AlphaFold2 in order to predict the structures of proteins. After using this tool or service, the author(s) reviewed and edited the content as needed and take(s) full responsibility for the content of the publication. The authors used ChatGpt to generate icons that are used to illustrate the species in the studies, based on in-house artwork.

## SUPPLEMENTAL INFORMATION

**Document S1. Figures S1–S4 and Supplementary Items S1 and S2 related to Figure 1**

## STAR★METHODS

### KEY RESOURCES TABLE

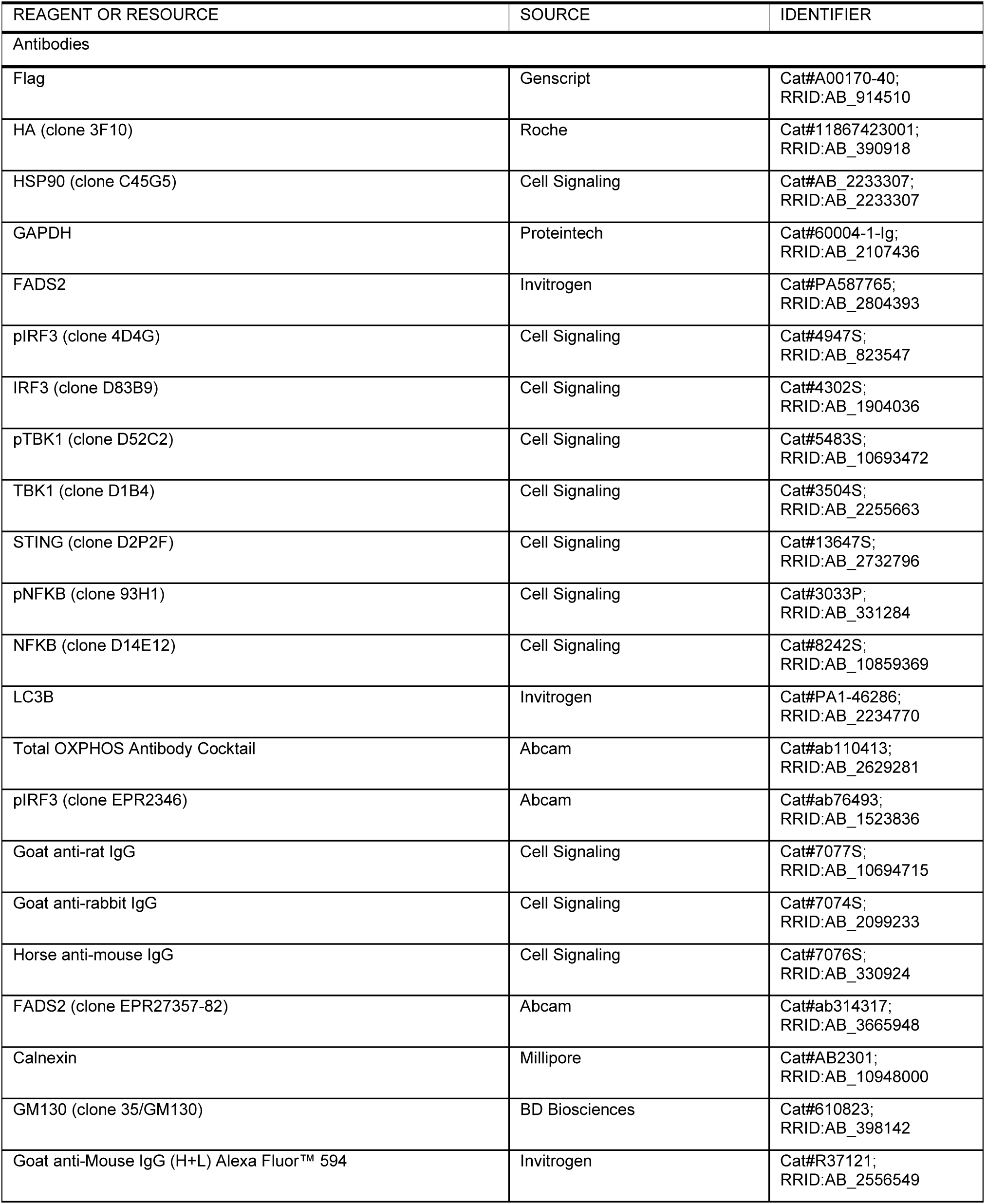
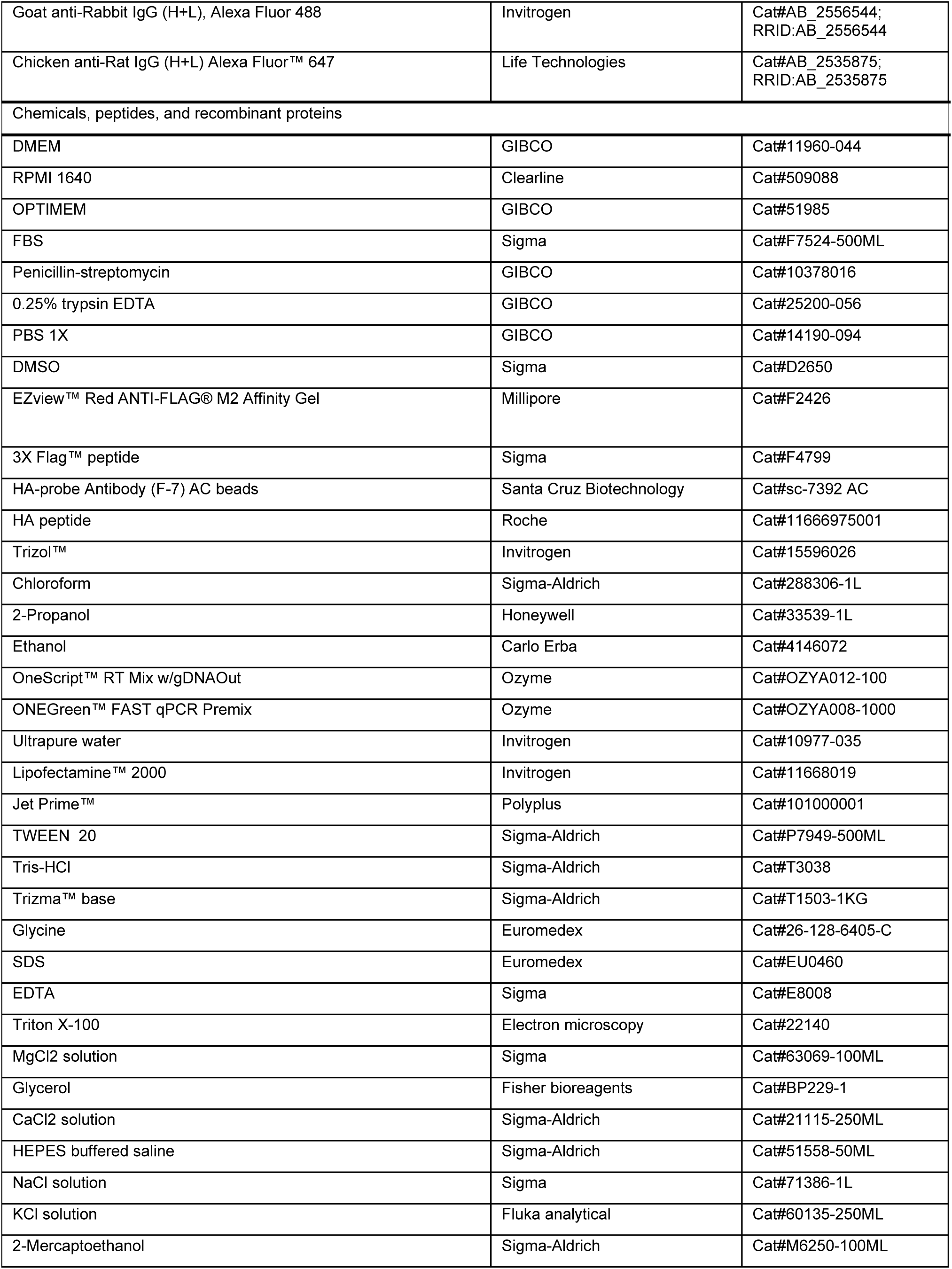
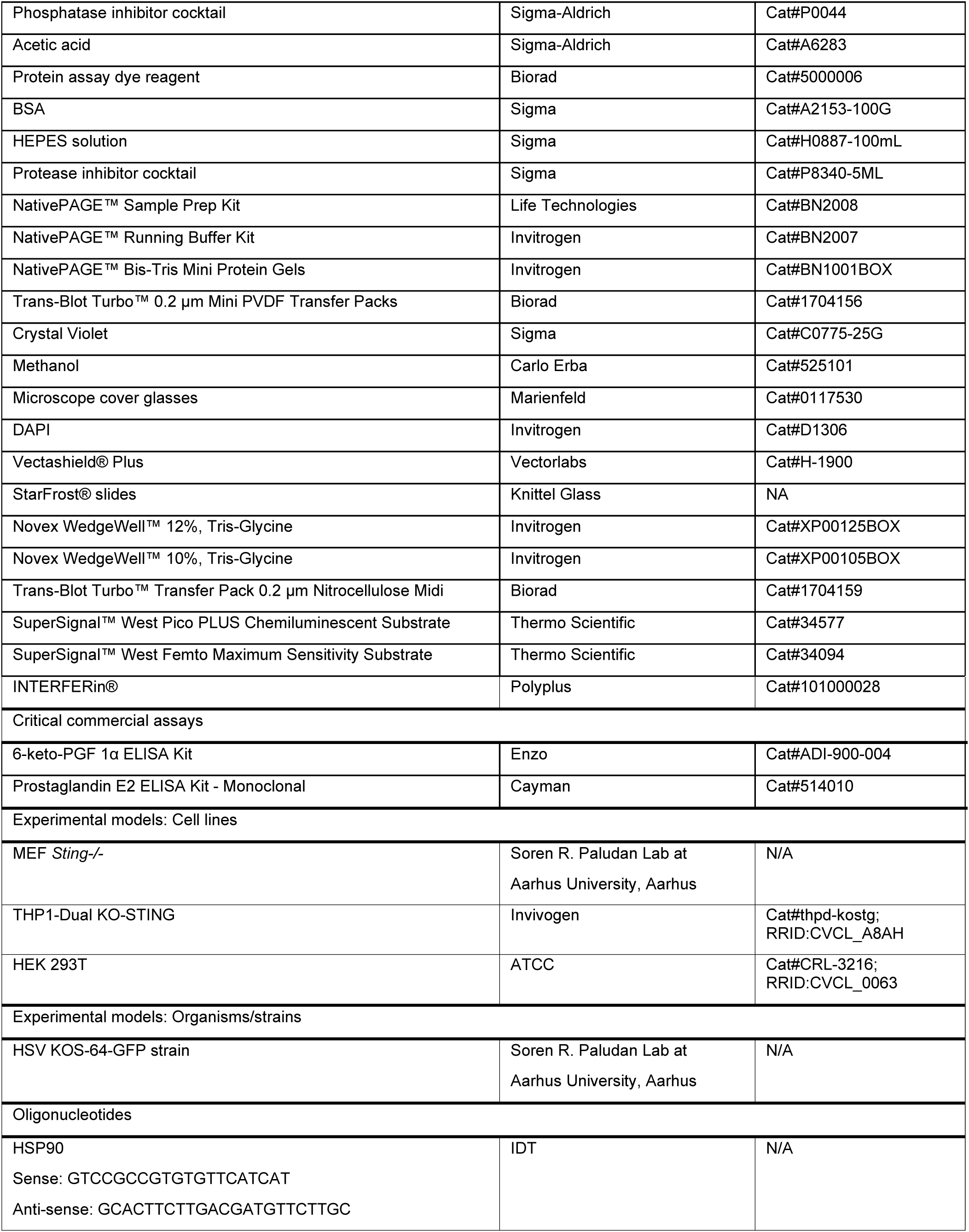
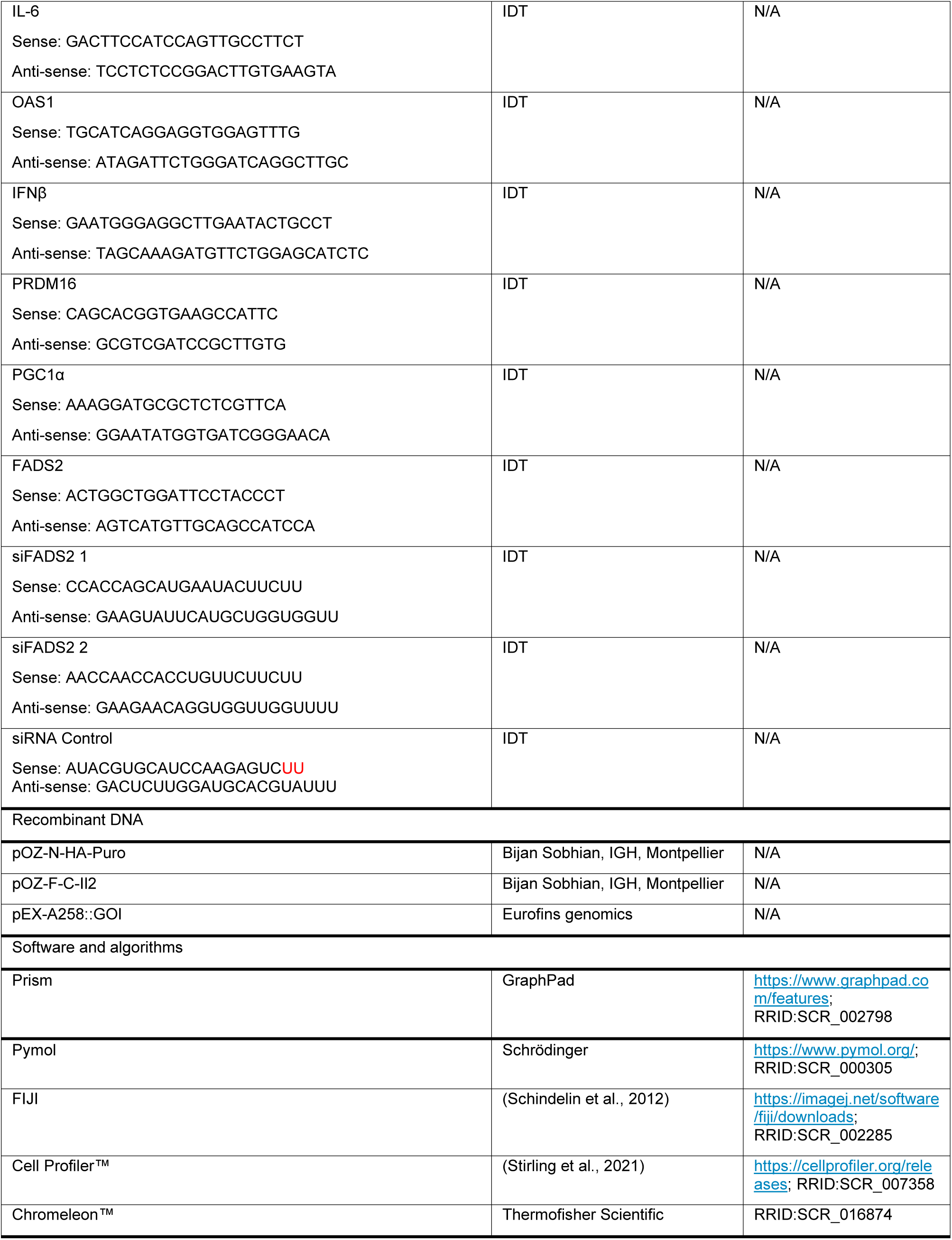

### EXPERIMENTAL MODEL AND STUDY PARTICIPANT DETAILS

Please list here under separate headings all the experimental models/study participants (animals, human participants, plants, microbe strains, cell lines, primary cell cultures) used in the study. For each model, provide information related to their species/strain, genotype, age/developmental stage, sex (and gender, ancestry, race, and ethnicity if reported for human studies), maintenance, and care, including institutional permission and oversight information for the experimental animal/human participant study. The influence (or association) of sex, gender, or both on the results of the study must be reported. In cases where it cannot, authors should discuss this as a limitation to their research’s generalizability.

### METHOD DETAILS

#### Cells and cell cultures

HEK 293T (ATCC), and MEF^Sting-/-^ (gift from Soren R. Paludan) were maintained in DMEM supplemented with 10% Fetal Bovine Serum (FBS), 100 µg/ml Penicillin-Streptomycin (Pen-Strep), and 1% L-Glutamine. THP1-Dual™ KO-STING (Invivogen) were maintained in RPMI 1640 w/ stable L-Glutamine supplemented with 10% FBS, 1% Penicillin-Streptomycin, and 100 µg/ml Normocin® at 5 x 10^5^ cells/ml. All cells were maintained at a temperature of 37 °C under 5% CO_2_. MEF^Sting-/-^ expressing N-terminal HA-tagged STING homologs/mutants (MbSTING, NvSTING, DrSTING, DrSTING-ΔCTT, XtSTING, MmSTING, and MmSTING-ΔCTT) and THP1-Dual™ KO-STING expressing N-terminal HA-tagged HsSTING haplotypes/mutants (HsSTING-R232H, HsSTING-AQ, HsSTING-HAQ, and HsSTING-R232H-ΔCTT) were generated by transduction of MEF*^Sting-/-^* or THP1-Dual™ KO-STING with retroviral particles packaging the corresponding pOZ-HA-STING or the pOZ-STING-HA constructs generated for each STING homolog/mutant/haplotype, followed by selection with 2 µg/ml puromycin.

#### Plasmid constructs

The pOZ-N-HA-Puro and pOZ-HA-C-Puro expression vectors used for expression of STING homologs and pOZ-FLAG-C-Il2 expression vector used for expression of FADS2 and FADS-like enzymes were obtained from Bijan Sobhian, IGH, Montpellier. The sequences corresponding to STING homologs (FsSTING, MbSTING, NvSTING, CgSTING, DrSTING, XtSTING, and MmSTING), the FADS2 homologs (MbFADS, NvFADS, DrFADS2, XtFADS2, and MmFADS2) were purchased from Eurofins Genomics as synthetic genes. The DrSTING-ΔCTT, and MmSTING-ΔCTT were generated in-house by deleting the C-terminal tail of the DrSTING and MmSTING. The sequence for HsSTING-R232H was also obtained from Bijan Sobhian, IGH, Montpellier and the AQ, HAQ, and ΔCTT of HsSTING were generated in-house from this sequence. The sequences corresponding to the STING and FADS homologs are:

FsSTING (Uniprot ID: P0DUD7):

MEKIYTWLKTNSYIVHHVSTSLNIISFIIVLIWIFESTIKEKLNIIFTVNLEAIVVFISILIVGLNQLLQKLLIEAEYSPAFALAVGYFKNFIFPAITQIKENGEVNPKICIYKPKHFDELTSTNIDMIKAELTNKKYNLSEINLSLKGARARDILTLNKKSKIHSYFDFPNTLLSLYSYVDFKIASSNNNSSELKKKKFVELLIEQFYLKLNELIQENNLTNNITF CDKNLQGL

MbSTING (Uniprot ID: A9UUE7)

MMVNLSDLSHLSQRGWAQVFVTLAAAAISIFYFSFSPFEVAAGICASIAAAGAVPLLFDAVHYLIAFCISAPDARPPLRTVWTKTRLQRWSGLSILTFIVLACGLYFASPSPMSASNLFAVFALSLCNSVFSAMVAPRCGRLEVVDLATHPNTVLFRDFGAASAMSYWHGYLQHIAGDRAALEERFRDSDVGGMRVPRKLYILVPQDCEVNASDNKYEGLKATEHYISPKPITIGGVVDREMGKHTLYTPSESAETASVAFAMELASPLNTMKNALKDAAEGLLELQSEAFYLTLRGILKQEGVLDQGNDEGMIKLVWGKNRSEVLKKMRAEDQYAMP

NvSTING (Uniprot ID: A7SLZ2)

MRRAEENNGFGTIPKRRNQHTPFYASIGMIVVIIVAFTSYHITSYGDDRNRAMRQYSFTFSLAYLAFLVGELLRRCCLFAEEYRHIETRYNGSLKKAIQTTFSFGHNNVLFVASLLFFVVFVASNDPNGSSSVIQGNSTAEPHTEMRQTSGWQGLWGQFIISALLTPLVVHLLGLRELSKVEESQLNEKENKNVADGLAWSYYFGYLKFVLPELEKQIEKTSKFRSKEKFVKKMFILIPSNCFWDDKIPGSDYDPQNRITFEGNTEPLEKTRGGVFLRHYKHSVYEIKDGENEPWFCIMEYATPLLTLYDMSVAQPGELSREERDAQVVVFLRKLQDILEGDRACQGKYELVTFSPDRDLADVMLRKLKDSELEIGG

CgSTING (P0DUE1):

MEKNGAHSFLSDTPVTSLTMSVPVLRHPHVYHAFISYCADADTSHARTILDSVESRGFTCCFAERDFLPGECTSDVVVDAIHCSKNVILVISPASLQSEWSKFEMLMAVDDSHQRNNVCLVPVLLGGVKVDDLPPPLRPLTCIELMDDFRNTDDIIQAISKPEDTWESLLPVGNLAHGFAWGYYYGYLKIILPDLDKTVRQWRRVNNAEGRMSEKLFLFFPQSCRCRDSIADESSLIKHRGHLPIITKDRAGIIERQYKNTIYSVTDDNGEDYFFAGEYIGVIHTMFEMEQNATTGLQTREKYVQSMRFYLTLKRILDTDPECSKKCKIVFYKDVNNSSDAMPKLICNEIKNQLRKESSDDTTVCMTPFNSPFPSISSPDFARCSLKSPSSTNMVKSEPNIYREESGKTKSVERG

DmSTING (A0A0B4LFY9):

MAIASNVVEAGNAVRAEKGRKYFYFRKMIGDYIDTSIRIVATVFLADLLLRLYRCVVEYGSNGRYYLPED RLWIILRRSCTYNNRSIYLIVGFLLVAFFRISVTGNYRNVMPTTLFLFQMPLYWIWSFTDMDQSTLSYSH WIRDSHGLDYAAGMASNYFHGYLKLSLPERKDDGLKHRLAMYEDKNNVTFGIKRLVILIPDEMFVNGVLE SHLLDKAEPLETQFINRAGVYRPFKHDVYRMNKKVNGRTYYFAVEGATPMISFFDATYSNLSGTWQMQE L KREIWIKFYKHLKELITTWPETRDLVELIIYNSHDSKGNLVDVGELLVAHMQNKTKTIDEISN

DrSTING (Uniprot ID:E7F4N7)

MSVMGEDALVPRARSRLPVMCAAGLGFLTLAVAWLLDSDKFSERAGIIAFGLMLERFIYCICLLAEELLFHSRQRYHGRMSEIFRACFRGSGILGMCAIFLMLMLGGVSFSVKQWSHFNLMCAGYMLLNSLGVLGPAPVEISEICEAKKMNVAHGLAWSFYIGYLKFLLPALEVNVREYSRRERLSSPRLHILLPLNARVPSKPEEEDTNVVFHENLPDLKLDRAGVRKRSYTNSVYKITHNNETFSCILEYATPLLTLYQMSQESSAGFGERERKQQVLLFYRTLSQILDNSLECRNRYRLILLNDEHTGDPHYLSRELFQNLKQQDGEIFMDPTNEVHPVPEEGPVGNCNGALQATFHEEPMSDEPTLMFSRPQSLRSEPVETTDYFNPSSAMKQN

XtSTING (Uniprot ID: A0A8J1J6F3)

MASIRNTLATQNRQIIPERRGKRATKMACVLAIGSILFVWILGKGKYSGAQLIYRMAINFAISQGCCLVTCACELTEEIKHLHTRYNGHYWRALKASFNLSCAAFVTAILCYVFYEPKLMASLPLTIDITLTLLSWLFCWILGIQGPTPATISEITEIKQLNVAHGLAWSYYVGYLQFVLPALKESIQKFNEENHNLLKFPETCRLHILIPLSCRLYGDLKDVDENITFLKEIPPLYIDRAGIKGRVFKNNVYRILDEDGRPYNCIVEYATPLASLLKMTDIPSAAFSADDRLQQTKLFYRTLKDILENAHELQNTYRLIVYEDFPETKDHSRHLLSQEILKHIRQQHSEEYSML

MmSTING (Uniprot ID: Q3TBT3)

MPYSNLHPAIPRPRGHRSKYVALIFLVASLMILWVAKDPPNHTLKYLALHLASHELGLLLKNLCCLAEELCHVQSRYQGSYWKAVRACLGCPIHCMAMILLSSYFYFLQNTADIYLSWMFGLLVLYKSLSMLLGLQSLTPAEVSAVCEEKKLNVAHGLAWSYYIGYLRLILPGLQARIRMFNQLHNNMLSGAGSRRLYILFPLDCGVPDNLSVVDPNIRFRDMLPQQNIDRAGIKNRVYSNSVYEILENGQPAGVCILEYATPLQTLFAMSQDAKAGFSREDRLEQAKLFCRTLEEILEDVPESRNNCRLIVYQEPTDGNSFSLSQEVLRHIRQEEKEEVTMNAPMTSVAPPPSVLSQEPRLLISGMDQPLPLRTDLI

MbFADS (Uniprot ID: A9UU07)

MKICIDQEWYDLTKWAKYHPGGVRILERFDNQDATDHFYSLHSTDAIRKFKAMRPTETKEDVPEVLPIDASFRELRAKLMDDGWWDRDLLTELGILIPIFAMCIIGTALAWSHPVISVLLISVGMQQAGWLGHDMTHARDSRYNDFWLRYVSGWLNGFDRNWWSNKHNTHHVLTNHVNHDPDIHVQPILYLWAPLKQMDHFLRKYQYIYFPLPYTLLFASWRLESLKWSIANRDYKMFFLAILPGYVWLALLPFKVVLASILVSGFLVAIVVTMSHESEELLLEREPSYVTNQFLTTRDVQCLDWVTEYLFGGMQYQLEHHLFPTMPRYKYRALVPIVRQWAKANGLVYKSDTLPVMLGEHVATLKKAGEMAHREDTVDPYALS

NvFADS (Uniprot ID: A7RJB6)

MTVALQRRTPKVYTLDEVKEHCSKGDCWVVVEDSVYDLSKWIGHHPGGELPILYMAGRECTDVFKAFHPAWVFTKKLPAFKIGKLDDTRKEKKETSLSEDFEKLRQEIIEGGGLQTNYWFYIRLASMLALLFASIIYCVVFSNNVYIQVAAGILVAVFWQQMAFIGHDAGHHAIFHDEQWDDRLGLVVGNLLTGVSIGWWKKSHNAHHVVTNSVELDPDIQHLPVLAVTDKFFNSIKSIYHDRVMHFDGLAKFFVRYQHHLYFLIMGLARFNLYAQSFLLVLSKERVKLRVMEFVTMVLFWTWYLTLCSYLPTWSTRFAFVFLAHFLAGIIHIQITLSHFSMETYNGLPLDAFKENRFLLSQMDTTMDIECDPNLDFFHGGLQFQFEHHLFPRVARQNLRSIQEKMKLLCKKHGLPYRSKSFVDANIEVIQCLKDTAEKSKCFSPKIWDSVHCIG

DrFADS2 (Uniprot ID: Q9DEX7)

MGGGGQQTDRITDTNGRFSSYTWEEVQKHTKHGDQWVVVERKVYNVSQWVKRHPGGLRILGHYAGEDATEAFTAFHPNLQLVRKYLKPLLIGELEASEPSQDRQKNAALVEDFRALRERLEAEGCFKTQPLFFALHLGHILLLEAIAFMMVWYFGTGWINTLIVAVILATAQSQAGWLQHDFGHLSVFKTSGMNHLVHKFVIGHLKGASAGWWNHRHFQHHAKPNIFKKDPDVNMLNAFVVGNVQPVEYGVKKIKHLPYNHQHKYFFFIGPPLLIPVYFQFQIFHNMISHGMWVDLLWCISYYVRYFLCYTQFYGVFWAIILFNFVRFMESHWFVWVTQMSHIPMNIDYEKNQDWLSMQLVATCNIEQSAFNDWFSGHLNFQIEHHLFPTVPRHNYWRAAPRVRALCEKYGVKYQEKTLYGAFADIIRSLEKSGELWLDAYLNK

XtFADS2 (Uniprot ID: A0A6I8QWG3)

MGMGGQSGEGCSSGNCVKPEARYSWEEIQKHNLKTDKWLVIERKVYNITQWVKCHPGGMRVIGHYAGEDATDAFHAFHPDKNFVRKFLKPLYVGELAENEPSQDRDKNAQQVEDFRALRKTAEDMGLFKSNPAFFIVYLFHILLIEFLAWCTLHYFGTGWIPTILTVLLLTISQAQAGWLQHDFGHLSVFKKSKWNHLIHKFIIGHLKGASANWWNHRHFQHHAKPNIFRKDPDVNMVNVFVLGDTQPVEFGKKRIKYLPYNHQHLYFFLIGPPLLIPIYFTIQIMKTMITRKDWVDLAWSVSYYARFFITFVPFYGVLGSLALLNAVRFIESHWFVWVTQMNHLPMVIDHEKYKDWLETQLAATCNIEPSFFNDWFSGHLNFQIEHHLFPTMPRHNYWKIAPLVRSLCSKYNVTYEEKCLYHGFRDVFRSLKKSGQLWLDAYLHK

MmFADS2 (Uniprot ID: Q9Z0R9)

MGKGGNQGEGSTERQAPMPTFRWEEIQKHNLRTDRWLVIDRKVYNVTKWSQRHPGGHRVIGHYSGEDATDAFRAFHLDLDFVGKFLKPLLIGELAPEEPSLDRGKSSQITEDFRALKKTAEDMNLFKTNHLFFFLLLSHIIVMESLAWFILSYFGTGWIPTLVTAFVLATSQAQAGWLQHDYGHLSVYKKSIWNHVVHKFVIGHLKGASANWWNHRHFQHHAKPNIFHKDPDIKSLHVFVLGEWQPLEYGKKKLKYLPYNHQHEYFFLIGPPLLIPMYFQYQIIMTMISRRDWVDLAWAISYYMRFFYTYIPFYGILGALVFLNFIRFLESHWFVWVTQMNHLVMEIDLDHYRDWFSSQLAATCNVEQSFFNDWFSGHLNFQIEHHLFPTMPRHNLHKIAPLVKSLCAKHGIEYQEKPLLRALIDIVSSLKKSGELWLDAYLHK.

#### RNA extraction and quantitative real-time PCR

Total RNA was extracted using TRIzol™ reagent (Invitrogen) from cell pellets. RNA was quantified with a NanoDrop™ spectrophotometer and 1 µg RNA was reverse transcribed using OneScript® RT Mix (Ozyme) and gDNAOut Mix (Ozyme). Gene expression analysis of specific genes was performed by using through real-time PCR using ONEGreen® FAST qPCR Premix (Ozyme) and appropriate primers in a CFX Opus 384 (Biorad). Reactions were performed in duplicates and relative quantification of target cDNA was done using the 2^-ΔΔCT^ method by normalizing to heat shock protein 90 (Hsp90) cDNA. Graphs were plotted and statistical analysis was done using Graphpad Prism.

The primer sequences used for RT-qPCR analyses are listed in the key resources table.

#### Cell seeding, Transfection, and treatments

For immunoprecipitation (IP) experiments with transiently overexpressed proteins, 4 x 10^6^ HEK 293T cells were seeded per 150 mm culture dish 24 hrs before transfection. For each dish, 12.5 µg of pOZ construct expressing each of the target protein was transfected using CaCl_2_. The transfection medium was replaced with fresh medium after 8 hrs and incubated for 48 hrs before harvest by scraping. For IP experiments with stably expressed proteins, 2 x 10^6^ MEF*^Sting-/-^* cells expressing the indicated STING homolog was plated per 150 mm culture dish and incubated for 24 hrs prior to harvest by scraping.

2×10^5^ MEF cells were seeded per well of 6-well plates 24 hrs before 2’3’-cGAMP treatment. Cells were treated with 4 μl Lipofectamine® 2000 reagent (Invitrogen) with or without 14 µM 2’3’-cGAMP in 1 ml total volume of Opti-MEM Reduced Serum Medium (Gibco) per well. For gene expression analyses by RT-qPCR, cells were harvested 9 hrs following 2’3’-cGAMP transfection. For protein detection and quantification by western blot, cells were harvested 6 hrs following 2’3’-cGAMP transfection. In western blot analyses where treatment with chloroquine was necessary to enhance LC3B-lipidation, cells were pre-treated or not with 20 µM chloroquine before 2’3’-cGAMP transfection in the presence or absence of 20 µM chloroquine. Analysis of STING oligomerization with native gel was done following 1 hr or 3 hrs of treatment with 2’3’-cGAMP. For immunofluorescence experiments, cells were seeded in poly-D-lysine coated coverslips followed by transfection of 2’3’-cGAMP for 3 hrs.

For transfection with siRNA, 10 nM siRNA duplex complexed with INTERFERin® transfection reagent (Polyplus) in Opti-MEM Reduced Serum Medium (Gibco) was used according to the manufacturer’s instructions to reverse transfect 0.7 x 10^6^ MEF*^Sting-/-^* cells expressing the indicated STING homolog plated per well of 6-well plate in 2 ml of antibiotic-free complete DMEM and incubated for 24 hrs. A second round of transfection with siRNA was performed following the same procedure and gene and protein expression analysis was performed 48 hrs later.

dsDNA probes used for the stimulation of STING were generated by annealing of sense and anti-sense ssDNA probes (listed in the key resources table) and integrity assessed as described in (Chamma et al., 2022). 2×10^5^ MEF cells were seeded per well of 6-well plates 24 hrs before transfection. 2 µg of dsDNA probes were transfected per well using 4 ul of JetPrime® transfection reagent (Polyplus), in 2 ml of complete DMEM. For ELISA, cells were harvested 24 hrs after transfection.

For plaque quantification assay after HSV-KOS64 infection, 5 x 10^4^ MEF cells were seeded per well of 24-well plates 24 hrs prior to infection.

#### Western blot analyses

Protein samples for denaturing western blots were prepared by lysing the cell pellet in 5 times the packed cell volume of TENTG-150 buffer (20 mM Tris-HCl pH 7.4, 0.5 mM EDTA, 150 mM NaCl, 10 mM KCl, 0.5% Triton X-100, 1.5 mM MgCl₂, and 10% glycerol, supplemented with 10 mM β-mercaptoethanol, 0.5 mM PMSF, and 1× phosphatase inhibitor) for 30 minutes at 4 °C in a rotary mixer. Lysate was then clarified by centrifugation at 14,000 × g for 30 minutes at 4 °C, and the resulting supernatant was stored at -80 °C. 20 µg of solubilized protein was mixed with Laemmli buffer and heated at 95 °C for 5 mins before separation by SDS-PAGE on a 10% or 12% or 10-20% Tris-glycine gel (Invitrogen Novex) and transferred to a nitrocellulose membrane (Bio-Rad Trans-Blot Turbo).

Protein samples for non-denaturing (native) western blots were prepared by lysing the cell pellet in native lysis buffer (25 mM NaCl, 20 mM HEPES pH 7.0, 10% glycerol, 1% DDM, supplemented with 1x protease inhibitor cocktail)(Chan et al., 2025) following the same procedure as mentioned above. Samples were prepared by adding the lysate to NativePAGE® sample buffer and NativePAGE™ 5% G-250 Sample Additive (Invitrogen). Samples were then run on a NativePAGE™ 3-12% Bis-Tris gel (Invitrogen) and transferred to a PVDF membrane (Bio-Rad Trans-Blot Turbo).

The western blot membranes were blocked with 5% milk in phosphate buffered saline (PBS) with 0.1% Tween® (PBST) for 30 mins at RT and probed with the indicated primary antibodies. The primary antibodies used are : anti-Flag (1:2000, Genscript, A00170-40); anti-HA (1:1000, Roche, 11867423001); anti-HSP90 (1:1000, Cell Signaling, 4877T), anti-GAPDH (1:5000, Proteintech, 60004-1-Ig), anti-FADS2 (1:5000, Invitrogen, PA587765), anti-pIRF3 (1:500, Cell Signaling, 4947S) , anti-IRF3 (1:1000, Cell Signaling, 4302S), anti-pTBK1 (1:1000, Cell Signaling, 5483S), anti-TBK1 (1:1000, Cell Signaling, 3504S), anti-STING (1:1000, Cell Signaling, 13647S), anti-pNFKB (1:500, Cell Signaling, 3033P), anti-NFKB (1:1000, Cell Signaling, 8242S), anti-LC3B (1:1000, Invitrogen, PA1-46286), Total OXPHOS Antibody Cocktail (1:250, Abcam, ab110413), pIRF3 (1:1000, Abcam, ab76493).

Horseradish peroxidase (HRP)-conjugated secondary antibodies (Cell Signaling Technology) were used at 1:2000 dilution. Immunoreactive proteins were visualized by chemiluminescent detection using SuperSignal™ West Pico Plus or Femto reagents (Thermo Scientific). Secondary antibodies used in the study are: Goat anti-rat IgG (1:2000, Cell Signaling, 7077S), Goat anti-rabbit IgG (1:2000, Cell Signaling, 7074S), Horse anti-mouse IgG (1:2000, Cell Signaling, 7076S).

#### Immunofluorescence

Cells attached to coverslips were fixed with cold methanol for 5 mins at -20 °C and blocked with 5% bovine serum albumin (BSA) in PBST for 30 mins at RT. Coverslips were incubated with the indicated primary antibodies diluted in the blocking solution at 37 °C for 45 min.

Primary antibodies used are for immunofluorescence are: anti-FADS2 (1:50, Abcam, ab314317), anti-HA (1:100, Roche, 11867423001), anti-Calnexin (1:250, Millipore, AB2301), and anti-GM130 (1:250, BD Biosciences, 610823).

Next, coverslips were incubated at 37 °C for 45 min with the following secondary antibodies: Alexa Fluor 488-coupled goat anti-Rabbit IgG (Invitrogen, R37116), Alexa Fluor 594-coupled goat anti-Mouse IgG (Invitrogen, R37121), and Alexa Fluor 647-coupled chicken anti-Rat IgG (Invitrogen, A-21472), diluted in the blocking solution. Cells were then stained with DAPI by incubating in 1 µg/ml DAPI diluted in PBS for 2 mins at RT. Finally, coverslips were mounted in Vectashield antifade mounting medium. Images were acquired on the Zeiss confocal LSM880 Airyscan microscope using the Zeiss Zen software. Images analyzed using the FIJI image processing package. Protein colocalization analysis was performed with Cell Profiler. Graphs were plotted and statistical analysis was done using GraphPad Prism.

#### Immunoprecipitation

Cells were lysed in 5 times the packed cell volume of TENTG-150 buffer and the input of the immunoprecipitation experiments were prepared as mentioned in the ‘Western blot analyses’ section. The HA/FLAG-tagged FADS/STING was immunoprecipitated using either agarose beads conjugated to anti-HA antibody (Santa Cruz) or EZview™ Red anti-FLAG® M2 beads (Sigma-Aldrich). Lysate containing approx. 7 mg of total protein was incubated with 10 µl packed bead volume of appropriate beads for 4 hrs at 4 °C in a rotary mixer. The immunoprecipitated proteins along with their binding partners were eluted from the beads using an elution buffer containing either 1 mg/ml HA-peptide (Roche) or 200 ng/µl 3xFLAG™ peptide (Sigma-Aldrich) in TENTG-150. The eluate was then boiled in Laemmli buffer and analyzed by western blot.

#### HSV-KOS64 infection

HSV-KOS64 molecular clone expressing a GFP was a gift from Soren Paludan. Amplification and titration of HSV-KOS64 is described in(Vila et al., 2022).

MEF expressing the indicated STING homologs were infected with an MOI dilution series of HSV KOS-64-GFP in DMEM for 90 minutes and then the infection medium was replaced with DMEM containing 2% human serum. Sixteen hrs post-infection, the medium was changed to DMEM supplemented with 10% FBS and incubated for another 32 hrs. Cells were then fixed with 4% paraformaldehyde (PFA) for 10 mins at RT and stained with crystal violet. The number of plaques per condition were then counted. Graphs were plotted and statistical analysis was done using Graphpad Prism.

#### Prostaglandin ELISA

Intracellular PGE2 was extracted from the cells. Cells were pelleted and resuspended in 0.1 M potassium phosphate buffer (pH 7.4), then sonicated in the same buffer. The lysates were subsequently centrifuged at maximum speed for 30 minutes at 4 °C. The total concentration of PGE2 was determined in the supernatant using the Prostaglandin E2 ELISA kit (Cayman item 514010, MI, USA) and the 6-keto-PGF 1α EISA kit (Enzo Life Sciences, NY, USA) following the manufacturer’s instructions and the concentrations were normalized to the total protein. Graphs were plotted and statistical analysis was done using Graphpad Prism.

#### Identification and analysis of STING and FADS sequences

The identification of a complete array of STING and FADS homologs across species was conducted using several strategies in parallel. Sequences previously reported were used to conduct searches using TBLASTN or BLASTP against datasets available in NCBI or Ensembl. Automatic annotations of the different genomes were also searched. Protein sequences for all genes analyzed here were collected from GenBank or Ensembl. All accession numbers are provided in the Plasmid constructs subsection. Sequence analysis and protein structure analysis was determined using SMART (http://smart.embl.de/). Additional analysis was performed using InterProScan (EBI) when relevant. Protein sequence alignments and similarity computing were conducted using Clustal Omega (EBI Web Services). Synteny analysis was performed using Genomicus (https://www.genomicus.bio.ens.psl.eu/genomicus-109.01/cgi-bin/search.pl), or NCBI or Ensembl, through extraction of relevant gene location from genome data viewers.

#### Lipidomics

The fatty acid composition of cells was determined according to previously published procedures(Carré et al., 2023; Lapaquette et al., 2024). Total lipids were extracted from cell pellets by using the Folch’s procedure(Folch et al., 1957). Transmethylation of fatty acids and fatty alcohols into fatty acid methyl esters (FAMEs) and dimethylacetals (DMAs), respectively, was achieved by using boron trifluoride in methanol according to Morrison and Smith(Morrison and Smith, 1964). FAMEs and DMAs were then extracted with hexane. The composition of FAMEs and DMAs was determined by gas chromatography coupled to flame ionization detection on a GC Trace 1310 (Thermo Scientific, Les Ulis, France) gas chromatograph using a CPSIL-88 column (100×0.25mm i.d., film thickness 0.20 μm; Varian, Le Plessis-Robinson, France). Hydrogen at 200 kPa was used as carried gas. The oven temperature was set at 60°C for 5 min, then increased to 165 °C at 15 °C/min, hold at 165 °C for 1 min, increased to 225 °C at 2 °C/min, and then hold at 225 °C for 17min. The injector and the detector were maintained at 250 °C. Individual species of FAMEs and DMAs were identified by comparison to commercial and synthetic standards. Chromeleon 7 software (Thermo Scientific) was used to determine the relative quantities of FAMEs and DMAs. Graphs were plotted and statistical analysis was done using Graphpad Prism.

### QUANTIFICATION AND STATISTICAL ANALYSIS

Statistical analysis was performed using GraphPad Prism. To compare normally distributed data from two groups/conditions, p-values were determined by unpaired two-tailed Student’s t-test followed by Holm-Sidak multiple corrections test was performed. For non-normal data, p-values were determined by Mann-Whitney test. All analyses and plots were generated from at least 3 independent biological replicates. The exact number of replicates and statistical analysis used for each experiment are indicated in the figure legends. All data were expressed as mean ± SD. p-value < 0.05 was considered significant (*), p < 0.01 (**), p < 0.001 (***).

