## Supplementary Figures for "Regulation of lipid metabolism is a primordial function of STING"

**SUPPLEMENTARY FILE**

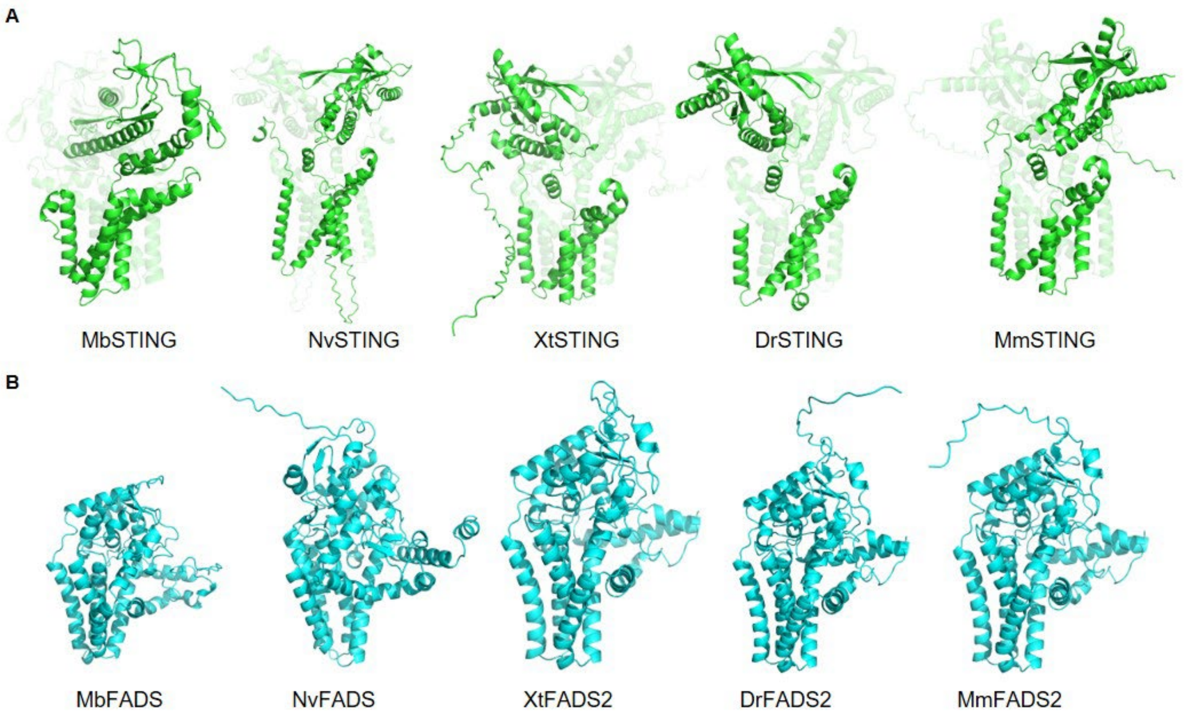


***Supplementary figure 1. Comparative analyses of STING and FADS homologs.***

(A) 3D Structures of STING in the indicated species as predicted by Alphafold2 were rendered using Pymol.
(B) 3D Structures of FADS2/FADS-like enzymes in the indicated species as predicted by Alphafold2 were rendered using Pymol.
Related to Figure 1

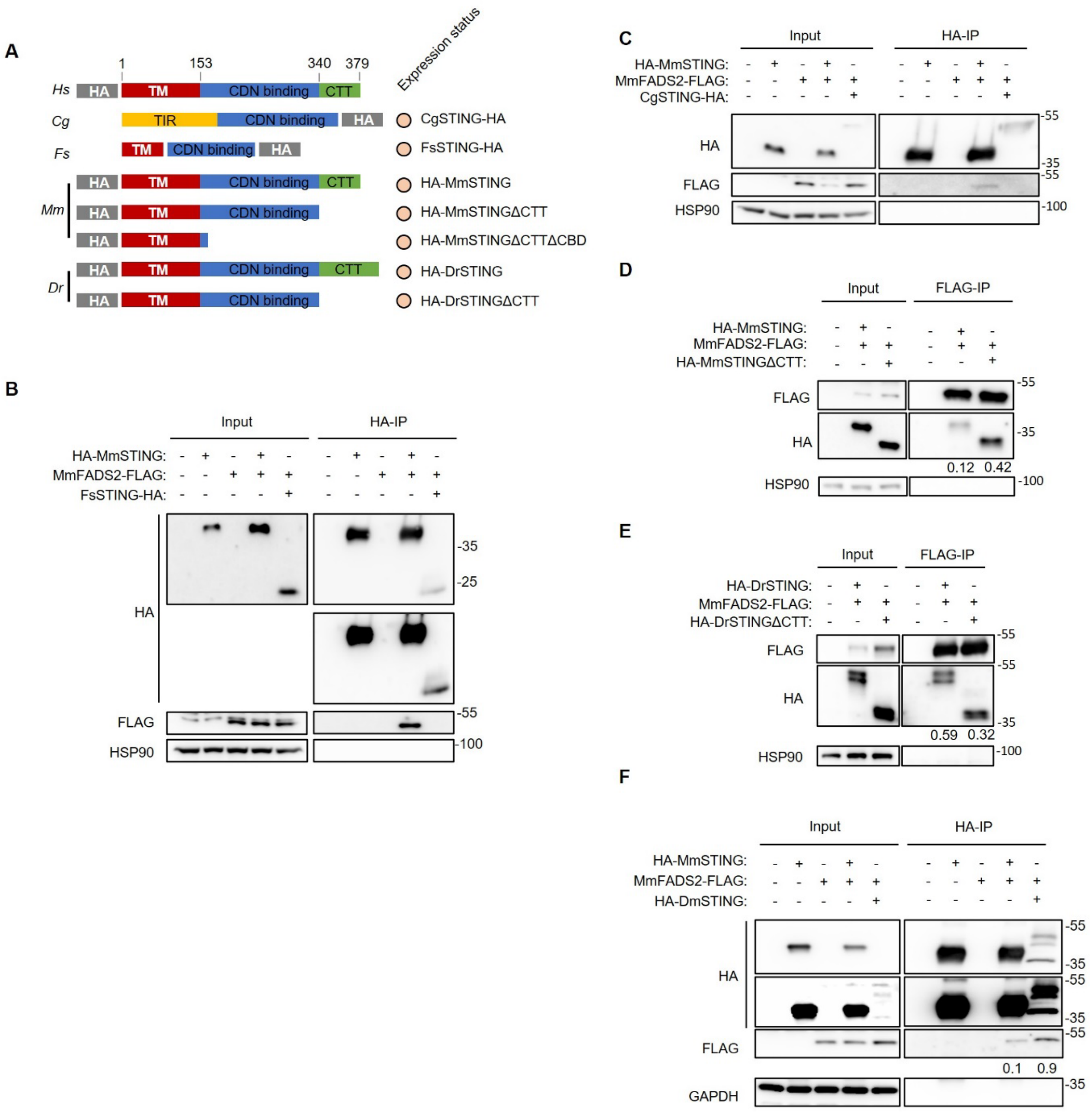


***Supplementary figure 2. The CTT is dispensable for interaction of STING with FADS2.***

(A) Schematics of HA-tagged constructs of STING with rare domains or STINGΔCTT mutants roughly aligned to HsSTING and their expression status in HEK 293T cells indicated (orange: present, white: absent).
(B) HA-immunoprecipitation was performed on WCEs from HEK 293T cells transiently co-expressing MmFADS2-FLAG or not with FsSTING-HA, HA-MmSTING (positive control), or none. Inputs and eluates from the immunoprecipitation were analyzed by WB using the indicated antibodies.
(C) As in B, except that cells expressed CgSTING-HA.
(D) FLAG-immunoprecipitation was performed on WCEs from HEK 293T cells transiently co-expressing MmFADS2-FLAG or not with MmSTINGΔCTT, HA-MmSTING (positive control), or none. Inputs and eluates from the immunoprecipitation were analyzed by WB using the indicated antibodies.
(E) As in D, except that cells expressed HA-DrSTINGΔCTT.
(F) As in B, except that cells expressed HA-DmSTING.
WBs are representative of 3 independent experiments. The normalized binding index (indicated below the co-immunoprecipitated protein) for the immunoprecipitation experiments was calculated by normalizing the ratio between band intensities of the immunoprecipitated prey and bait to that of the prey input.
Related to Figure 2

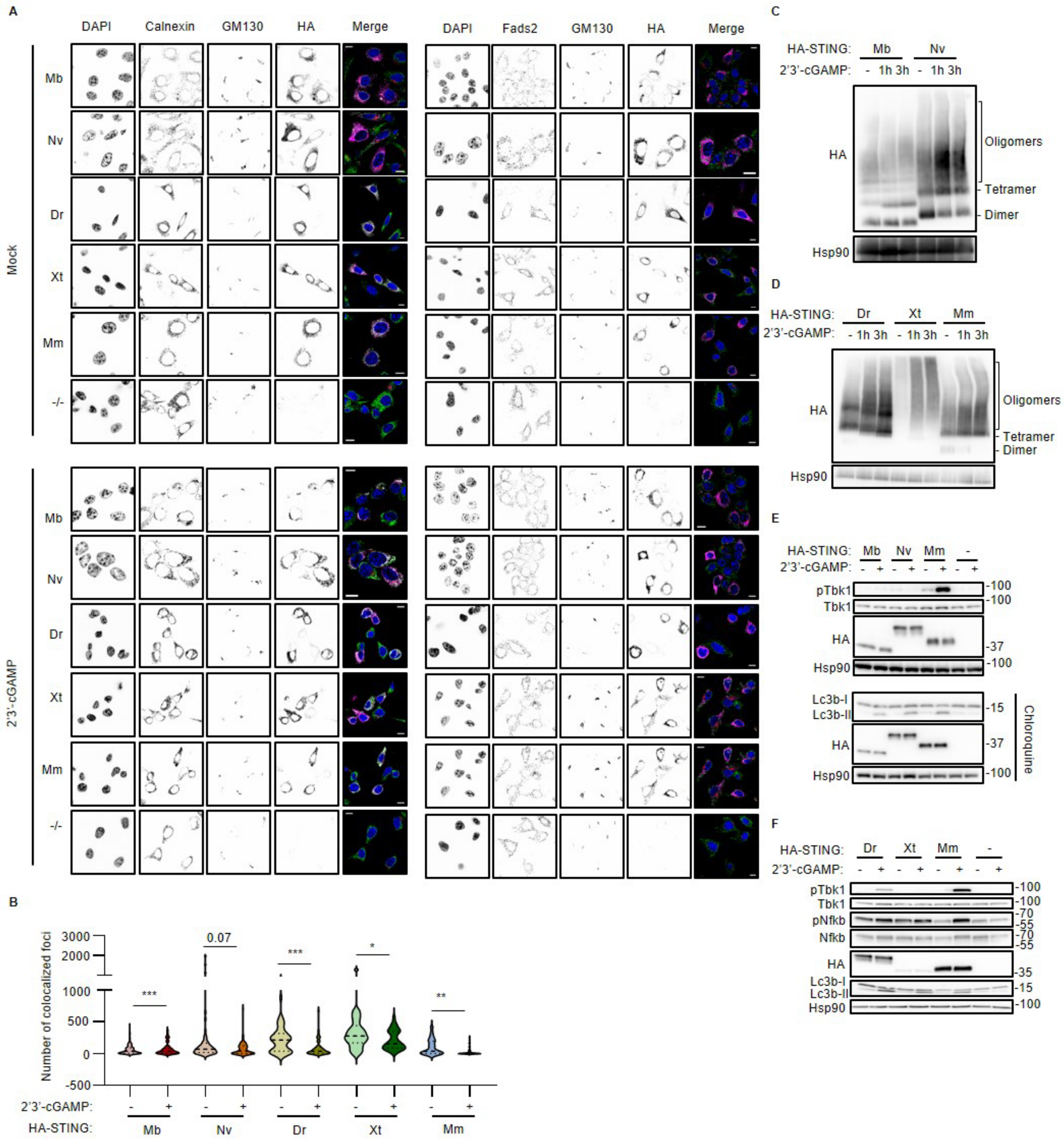


*Figure 3. The STING homologs are functional in MEF$^{Sting-/-}$ cells.*

(A) Immunofluorescence analysis of MEF$^{Sting-/-}$ stably expressing HA-STING homolog from the indicated species or parental cells, with or without stimulation with 14 µM 2'3'-cGAMP for 3h, using antibodies against the indicated proteins along with DAPI nuclear staining. Scale bar: 10µm. Images are representative of 3 independent experiments.
(B) Beanplot of the number of STING-Calnexin colocalization foci with or without 2'3'-cGAMP treatment. Mann-Whitney test was used to calculate p-values.
(C) Oligomerization status of the indicated vertebrate STING homologs in MEF$^{Sting-/-}$ treated or not with 14 µM 2'3'-cGAMP for the indicated durations analyzed by Blue-Native PAGE of WCE followed by WB using the indicated antibodies.
(D) As in C, except that the indicated invertebrate STING homologs were analyzed.
(E) WCEs from the same cell lines as in C, stimulated or not with 14 µM 2'3'-cGAMP for 6h and compared to parental MEF$^{Sting-/-}$ cells and analyzed by WB using the indicated antibodies.
F) As in E, except that the indicated invertebrate STING homologs were analyzed and 2'3'-cGAMP treatment was performed in the presence of chloroquine after 3h of pre-treatment with 20 µM chloroquine (bottom panel only). WBs are representative of 3 independent experiments.
Related to Figure 3

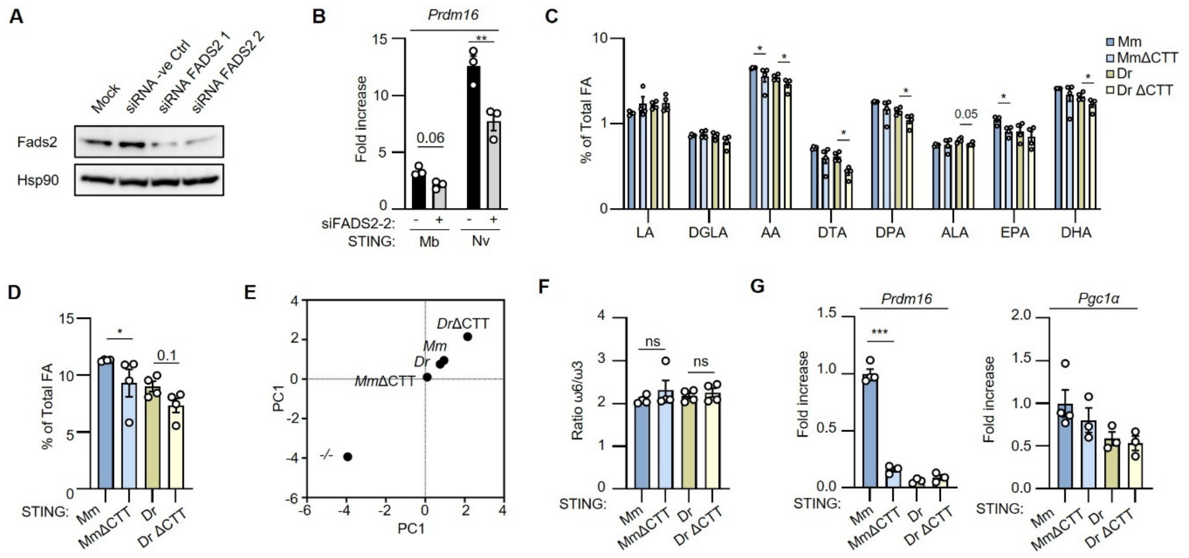


***Supplementary Figure 4. The CTT of vertebrate STING is dispensable for FADS2 activoty regulation***

(A) MEF WT cells were treated with 10 nM control or siFADS2 (siFads2-1 and siFads2-2), prior to WB analysis using the indicated antibodies. WB is representative of 2 independent experiments;
(B) RT-qPCR analysis of *Prdm16* levels was performed on MEF treated with control or siFADS2-2. Graph presents mean (±SEM) of n=3.
(C) Relative quantities of the indicated PUFAs and LC-PUFAs in MEF$^{Sting-/-}$ cells expressing the indicated HA-STING or HA-STINGΔCTT mutant measured using LC-MS and expressed as percentage of total fatty acids. Graph presents the mean (±SEM) of n=4.
(D) Summation of the LC-PUFA products of FADS2 measured in C, expressed as percentage of total fatty acids. Graph presents the mean (±SEM) of n=4.
(E) PC score plot from principal component analysis performed using percentages of FADS2 products as variables using the data from C.
(F) Ratio of total omega-6 to omega-3 PUFAs as measured in C. Graph presents the mean (±SEM) of n=4.
(G) RT-qPCR analysis of *Pgc1α* and *Prdm16* mRNA levels in the cell lines from C, normalized to *Hsp90* and expressed as fold increase over MEF$^{Sting-/-}$. Graph presents the mean (±SEM) of n=3-4.
Related to Figure 3.
