## Supplementary Items for "Regulation of lipid metabolism is a primordial function of STING"

```
HOMSA      -------MPHSSLHPSIPCPRGHGAQKAALVLLSACLVTLW-------GLGEPP-EHTLR	45
XENTR      -MASIRNTLATQNRQIIPERRGKRATKMACVLAIGSILFVW-------ILGKGK-YSGAQ	51
DANRE      -------MSVMGEDALVPRARSRLPVMCAAGLGFLTLAVAW-------LLDSDK-FSERA	45
DROME      -MAIASNVVEAGNAV--RAEK-------------GRKYFYF----R-KMIGDYIDTSIRI	39
CRAGI      -------MEKNGAHSFLSDTPVTSLTMSVPVLRHPHVYHAFISY------CADADT----	43 TIR
NEMVE      ----MRRAEENNGFGTIPKRRNQHTPFYASIGMIVVIIVAFTSYHI-TSYGDDRNRAMRQ	55
MONBE      MMVNLSDLSH------LSQRGWAQVFVTLAAAAISIFYFSFSPFEVAAGICASIAAAGAV	54
                                                   :

HOMSA      --Y-LVLHLASLQLGLLLNGVCSLAEEL----RHIHSRYRGSYWRTVRA--CLGCP----	92
XENTR      LIYRMAINFAISQGCCLVTCACELTEEI----KHLHTRYNGHYWRALKA--SFNLS----	101
DANRE      GIIAFG-----LMLERFIYCICLLAEELL---FHSRQRYHGRMSEIFRA--CFRGS----	91
DROME      ----VATVFLADLLLRLYRCVVEYGSNG------RYYLPEDRLWIILRRSCTYNNR----	85
CRAGI      --------------------------------SHARTILDSVESRGFTC--CFAERDFLP	69
NEMVE      YSFTFSLAYLAFLVGELLRRCCLFAEEY----RHIETRYNGSLKKAIQT--TFSFG----	105
MONBE      --------PLLFDAVHYLIAFCISAPDARPPLRTVWTKTRLQRWSGLSI-----------	95
                                                         .

HOMSA      --L--RRGALLLLSIYFYYS--LPNA--------------------VGPPFTW-MLAL-L	124
XENTR      --C--AAFVTAI-LCYVFYE--PKLM--------------------ASLPLTI-DITL-T	132
DANRE      --G--ILGMCAIFLMLMLGG--VSFS--------------------V---KQWSHFNL-M	121
DROME      ----SIYLIVGF-LLVAFFRISVT-------GNYRNVMPTTLFLFQMP------------	121
CRAGI      GECTSDVVVDAI-HC---------------------SKNVILVISPASLQSEWSKFEMLM	107
NEMVE      --HNNVLFVASLLFFVVFVASNDPNGSSSVIQGNSTAEPHTEMRQTSGWQGLWGQFIISA	163
MONBE      ------L-TFIVLACGLYFASPSPMS----------------------ASNLFAVFALSL	126
                      .

HOMSA      -----------GLSQALNILLGLKG--------------------------LAPAEISAV	147
XENTR      -----------LLSWLFCWILGIQG--------------------------PTPATISEI	155
DANRE      -----------CAGYMLLNSLGVLG--------------------------PAPVEISEI	144
DROME      ----------------LYWIWSFTD-----------------------MDQSTLSYSHWI	142
CRAGI      AVDDSHQRNNVCLVPVLLGGVKVDDLPPPLRPLTCIELMDDFRNTDDIIQAISKPEDTWE	167
NEMVE      -----------LLTPLVVHLLGLRE--------------------------LSKVEESQL	186
MONBE      CNSV-----FSAMVAPRCGRLEVVD-------------------------LATH-----P	151
                                 .                             :

HOMSA      CEKGNFNVAHGLAWSYYIGYLRLILPELQARI-----RTYNQHYNNLLRGAVSQRLYILL	202
XENTR      TEIKQLNVAHGLAWSYYVGYLQFVLPALKESI-----QKFNEENHNLLKFPETCRLHILI	210
DANRE      CEAKKMNVAHGLAWSFYIGYLKFLLPALEVNV-----REYSRRE-----RLSSPRLHILL	194
DROME      RDSHGLDYAAGMASNYFHGYLKLSLPERKDDGLKHRLAMYEDKNN---VTFGIKRLVILI	199
CRAGI      SLLPVGNLAHGFAWGYYYGYLKIILPDLDKTV-----RQWRRVNN--AEGRMSEKLFLFF	220
NEMVE      NEKENKNVADGLAWSYYFGYLKFVLPELEKQI-----EKTSKFRS---KEKFVKKMFILI	238
MONBE      NTVLFRDFGAASAMSYWHGYLQHIAGDRAALE-----ERFRDSD--VGGMRVPRKLYILV	204
                 : . . * .:: ***:                                :: ::.

HOMSA      PLDCGV----PDNLSMADPNIRFLDKLPQQTGDRAGIKDRVY-SNSIYELLE-NGQRAGT	256
XENTR      PLSCRL----YGDLKDVDENITFLKEIPPLYIDRAGIKGRVF-KNNVYRILD-EDGRPYN	264
DANRE      PLNARV----PSKPEEEDTNVVFHENLPDLKLDRAGVRKRSY-TNSVYKITH-N-NETFS	247
DROME      PDEMFVNGVLESHL------LDKAEPLETQFINRAGVY-RPF-KHDVYRMNKKVNGRTYY	251
CRAGI      PQSCRC----RDSIADESSLIKHRGHLPIITKDRAGIIERQY-KNTIYSVTD-DNGEDYF	274
NEMVE      PSNCFWDDKIPGSDYDPQNRITFEGNTEPLEKTRGGVFLRHY-KHSVYEIKD-GENEPWF	296
MONBE      PQDCEVNA----SDNKYEGLKATEHYISPKPITIGGVVDREMGKHTLYTPSESAETASVA	260
           * .                               .*:  *   .: :*   .

HOMSA      CVLEYATPLQTLFAMSQYSQAGFSR--EDRLEQAKLFCRTLEDILADAPESQN-----NC	309
XENTR      CIVEYATPLASLLKMTDIPSAAFSA--DDRLQQTKLFYRTLKDILENAHELQN-----TY	317
DANRE      CILEYATPLLTLYQMSQESSAGFGE--RERKQQVLLFYRTLSQILDNSLECRN-----RY	300
DROME      FAVEGATPMISFFDATYSNLSGTWQMQELKREIWIKFYKHLKELITTWPETRD-----LV	306
CRAGI      FAGEYIGVIHTMFEMEQNATTGLQT--REKYVQSMRFYLTLKRILDTDPECSK-----KC	327
NEMVE      CIMEYATPLLTLYDMSVAQPGELSR--EERDAQVVVFLRKLQDILEGDRACQG-----KY	349
MONBE      FAMELASPLNTMKNALKDAAEGLLE------LQSEAFYLTLRGILKQEGVLDQGNDEGMI	314
              *    : ::                        *   *  ::
HOMSA      RLIAYQEP---ADDSSFSLSQEVLRHLRQEEKEEVTVGSL-KTSAVPSTS----------	355
XENTR      RLIVYEDFPETKDHSRHLLSQEILKHIRQQHSEEYSML----------------------	355
DANRE      RLILLNDE---HTGDPHYLSRELFQNLKQQDGEIFMDPTN-EVHPVPEEGPVGNCNGALQ	356
DROME      ELIIYNSHDSKGNLV--DVGELLVAHMQNKTKT------------IDEISN---------	343
CRAGI      KIVFYKDVNNSSDAMPKLICNEIKNQLRKESSDDTTVCMTPFNSPFPSISSPDFARCSLK	387
NEMVE      ELVTFSPDRDLADVMLRKLKDSELEIGG--------------------------------	377
MONBE      KLVWGKNRSEVLKKMRAEDQYAMP------------------------------------	338


HOMSA      ------TMSQEPELLISGMEKPLPLRTDFS---------------	379
XENTR      ---------------------------------------------	355
DANRE      ATFHEEPMSDEPTLMFSR---PQSLRSEPVETTDYFNPSSAMKQN	398
DROME      ---------------------------------------------	343
CRAGI      SPSSTNMVKSEPNIYREESGKTKSVERG-----------------	415
NEMVE      ---------------------------------------------	377
MONBE      ---------------------------------------------	338
```

**Supplementary Item 1.** Alignment of of STING ortholog sequences from indicated species. Residues highlighted in yellow: TM, purple: TIR, grey: CDN-binding, green: CTT. Sequences were aligned with CLUSTAL O(1.2.4) multiple sequence alignment tool.

```
HOMSA      MGKGGNQ----GEGAAEREVS--VPTFSWEEIQKHNLRTDRWLVIDRKVYNITKWSIQHP	54
XENTR      MGMGGQS----GEGCSSGNCVKPEARYSWEEIQKHNLKTDKWLVIERKVYNITQWVKCHP	56
DANRE      MGGGGQQ----TDRITDT--NGRFSSYTWEEVQKHTKHGDQWVVVERKVYNVSQWVKRHP	54
DROME      MVIEEWKKSGIATKFPTYRNSALITTHSWQKGKRQDDGAEGLWRINDGIYDFTSFIDKHP	60
CRAGI      MKPGPSKEMG-KEVYSDVSMAKENKHYTVEEVKRHNERDDNWLMINGKVYDLSRWAKKHP	59
NEMVE      --------------MTVALQRRTPKVYTLDEVKEHCSKGDCWVVVEDSVYDLSKWIGHHP	46
MONBE      ----------------------------------------MKICIDQEWYDLTKWAKYHP	20
                                                       ::   *:.: :   **

HOMSA      GGQRVIGHYAGEDATDAFRAFHPDLEFVGKFLKPLLIGELAPEEPS----QDHGKNSKIT	110
XENTR      GGMRVIGHYAGEDATDAFHAFHPDKNFVRKFLKPLYVGELAENEPS----QDRDKNAQQV	112
DANRE      GGLRILGHYAGEDATEAFTAFHPNLQLVRKYLKPLLIGELEASEPS----QDRQKNAALV	110
DROME      GGPFWIRETKGTDITEAFEAHHLTTAPEKM-IAKYKVRDAAEPRIYTLTLEEGGFYKTLK	119
CRAGI      GGAKILGHYAGEDATDAWTAFHNDKGYVSKFMKALYIGDVEDWDQ---------NETEIK	110
NEMVE      GGELPILYMAGRECTDVFKAFHPAWVFTKK-LPAFKIGKLDDT-------RKEKKETSLS	98
MONBE      GGVRILERFDNQDATDHFYSLHSTDAIR-KF------KAMRPTETK----EDVPEVLPID	69
           **   :    . : *: : : *

HOMSA      EDFRALRKTAEDMNLFKTNHVFFLLLLAH-IIALESIAWFTVFYFGNGWIPTLITAFVLA	169
XENTR      EDFRALRKTAEDMGLFKSNPAFFIVYLFH-ILLIEFLAWCTLHYFGTGWIPTILTVLLLT	171
DANRE      EDFRALRERLEAEGCFKTQPLFFALHLGH-ILLLEAIAFMMVWYFGTGWINTLIVAVILA	169
DROME      ERVREQLKTIDKRPKKKSDLI-------HLGLVVSLYL---LG-IASAKYNSLLALVLAS	168
CRAGI      KDFRKLRKYVEESGLMKVNPLFYVLHLLH-ILALEVLAYLVVWHWEGSMAAFFVAAVLLA	169
NEMVE      EDFEKLRQEIIEGGGLQTNYWFYIRLASMLALLFASIIYCVVF--SNNVYIQVAAGILVA	156
MONBE      ASFRELRAKLMDDGWWDRDLL------TELGILIPIFAMCIIG-TALAWSHPVISVLLIS	122
             ..            . :            : .       :          .   .: :

HOMSA      TSQAQAGWLQHDYGHLSVYRKPKWNHLVHKFVIGHLKGASANWWNHRHFQHHAKPNIFHK	229
XENTR      ISQAQAGWLQHDFGHLSVFKKSKWNHLIHKFIIGHLKGASANWWNHRHFQHHAKPNIFRK	231
DANRE      TAQSQAGWLQHDFGHLSVFKTSGMNHLVHKFVIGHLKGASAGWWNHRHFQHHAKPNIFKK	229
DROME      VALCWTVIVSHNYFHR----RDNWQ----MYAFNLGMMNFAAWRVSHALSHHIYPNSYF-	219
CRAGI      TAQAQAGWSQHDYGHMSVFPSHKMNHIFHHIVIGHIKAASSHWWNFRHFQHHAKPNVIKK	229
NEMVE      VFWQQMAFIGHDAGHHAIFHDEQWDDRLGLVVGNLLTGVSIGWWKKSHNAHHVVTNSVEL	216
MONBE      VGMQQAGWLGHDMTHA---RDSRYNDFWLRYVSGWLNGFDRNWWSNKHNTHHVLTNHVNH	179
                     *:  *         :        .        *       **   *

HOMSA      DPDVNMLHVFVLGEWQPIEY----GKK------KLKYLPYNHQHEYFFLIGPPLLIPMYF	279
XENTR      DPDVNMVNVFVLGDTQPVEF----GKK------RIKYLPYNHQHLYFFLIGPPLLIPIYF	281
DANRE      DPDVNMLNAFVVGNVQPVEY----GVK------KIKHLPYNHQHKYFFFIGPPLLIPVYF	279
DROME      DLELSMFEPLLCWVPNPHIK-----SKLMRYVSWV--TEPVAYALAFFIQMGTRIFYSLR	272
CRAGI      DPDVDIAFLFLIGDYIPKSW----GSK------KLGFMPYNFQHSYFFFLGPPLLLPIYF	279
NEMVE      DPDIQHLPVLAVTDKFFNSIKSIYHDRVMHFDGLAKFFVRYQHHLYFLIMGLAR-FNLYA	275
MONBE      DPDIHVQPILYLWA--PLKQ--------------MDHFLRKYQYIYFP-----LPYTLLF	218
           * ::     :                                   :*          :

HOMSA      QYQIIMTMIVHKNWVDLA----WAVSYYIRFFITYIPFYGILGALLFLN--FIRFLESH-	332
XENTR      TIQIMKTMITRKDWVDLA----WSVSYYARFFITFVPFYGVLGSLALLN--AVRFIESH-	334
DANRE      QFQIFHNMISHGMWVDLL----WCISYYVRYFLCYTQFYGVFWAIILFN--FVRFMESH-	332
DROME      HTN-------ILYWHDLLPL-----TIPIAIYLGTGGSLGIWICVRQWL--AMTSIASFS	318
CRAGI      HVENIFFVFKRRDYVDLL----MTVTFFYKFFWLYGPYYGGWGAFGLYM--FMRFLESH-	332
NEMVE      QSF--L-LVLSKERVKLRVMEFVTMVLFWTWYLTLCSYLPTWSTRFAFV--FLAHFLAG-	329
MONBE      ASWRLE----SLKWS-------IANRDYKMFFLAILPGYV-WLALLPFKVVLASILVSGF	266
                                          :                       : :

HOMSA      WFVWVTQMNHIVM------EIDQEAYRDWFSSQLTATCNVEQSF-FNDWFSGHLNFQIEH	385
XENTR      WFVWVTQMNHLPM------VIDHEKYKDWLETQLAATCNIEPSF-FNDWFSGHLNFQIEH	387
DANRE      WFVWVTQMSHIPM------NIDYEKNQDWLSMQLVATCNIEQSA-FNDWFSGHLNFQIEH	385
DROME      FCLIGLNAAHHDPEIYHEG-DANREDRDWGLFQVDTIIDRGDLKWSQFLVLTHFGDHVLH	377
CRAGI      WFTWVTQMSHLPM------RVEKEKEEDWFSLQLATTCNVDSSP-FNDWFTGHLNFQIEH	385
NEMVE      IIHIQITLSHFSMETYNGLPLDAFKENRFLLSQMDTTMDIECDP-NLDFFHGGLQFQFEH	388
MONBE      LVAIVVTMSHESE------ELLLEREPSYVTNQFLTTRDVQCLDWVTEYLFGGMQYQLEH	320
                    *                  :   *. :  :          .   :  :. *

HOMSA      HLFPTMPRHNLHKIAPLVKSLCAK-HGIEYQEKPLLRALLDIIRSLKKSGK--------L	436
XENTR      HLFPTMPRHNYWKIAPLVRSLCSK-YNVTYEEKCLYHGFRDVFRSLKKSGQ--------L	438
DANRE      HLFPTVPRHNYWRAAPRVRALCEK-YGVKYQEKTLYGAFADIIRSLEKSGE--------L	436
DROME      HLFPTLDHGLLPALYPVLYQTLDEFKGHLRECNHIEHMIGQHKQLLRIEPNPRAPGAGK-	436
CRAGI      HLFPTMPRHNLGKIAPLVKSLCKK-HNLPYRCSPLFTAFGDIVRMLKKSGE--------I	436
NEMVE      HLFPRVARQNLRSIQEKMKLLCKK-HGLPYRSKSFVDANIEVIQCLKDTAEKSKCFSPKI	447
MONBE      HLFPTMPRYKYRALVPIVRQWAKA-NGLVYKSDTLPVMLGEHVATLKKAGEMAHREDT--	377
           **** : :         :        .   . . :     :    *.   :

HOMSA      WLDAYLHK	444
XENTR      WLDAYLHK	446
DANRE      WLDAYLNK	444
DROME      --------	436
CRAGI      WYDAYYHG	444
NEMVE      WDSVHCIG	455
MONBE      -VDPYALS	384
```

**Supplementary Item 2.** Alignment of FADS2 and FADS-like enzyme sequences from indicated species. Residues in red: TM. Residues highlighted in yellow: Cyt b5-like heme/steroid binding domain, green: FAD domain. Sequences were aligned with CLUSTAL O(1.2.4) multiple sequence alignment tool.
